# Combined AMPK activation and ghrelin ameliorate cancer cachexia through complementary effects on energy homeostasis, inflammation, and wasting

**DOI:** 10.64898/2026.06.23.733859

**Authors:** V. González-Alvarez, S. Caamaño, A. Reimúndez, J. Cañas-Martín, A. Capelo-Diz, N. Seoane, A. Pensado-López, P. Benedikt, M. Schweiger, D. Viña, A. Vieites, F. Torres Andón, VM. Arce, R. Señarís

## Abstract

**Background:** Cancer-associated cachexia is characterized by progressive loss of skeletal muscle and adipose tissue driven by systemic inflammation and metabolic dysregulation. AMP-activated protein kinase (AMPK) is a central regulator of energy homeostasis, but its role in cachexia and its therapeutic potential remains incompletely defined. We investigated AMPK signaling during cachexia and whether pharmacological AMPK activation alone or combined with ghrelin could ameliorate disease manifestations.

**Methods:** Cachexia was induced in male C57BL/6 mice by Lewis lung carcinoma (LLC) implantation. Additional models included fibrosarcoma (CHX and MN/MCA1) and chronic lymphocytic choriomeningitis virus (LCMV) infection. AMPK was activated using AICAR and BC1618 (AB), alone or combined with ghrelin (AB+G). Metabolic, inflammatory, and functional outcomes were assessed in hypothalamus, skeletal muscle, adipose tissue, and serum.

**Results:** LLC-bearing mice developed cachexia characterized by reduced body weight, lean and fat mass, hypophagia, and elevated circulating IL-6 and corticosterone. Cachectic LLC mice displayed increased *Il6* and *Il1β* expression in hypothalamus, skeletal muscle, and white adipose tissue (WAT). Furthermore, AMPK activation failed to increase in hypothalamus or peripheral tissues despite profound energy deficit. A similar defect in AMPK responsiveness was observed in CHX and LCMV models, indicating a conserved feature of cachexia.

AB treatment in LLC mice reduced circulating IL-6 and corticosterone levels and decreased skeletal muscle atrogene expression and IL-6/STAT3 signaling, partially preserving muscle mass, fiber size, and grip strength. However, food intake remained low, and WAT was largely unresponsive, maintaining elevated *Il6* expression and tissue loss.

Ghrelin alone increased food intake in LLC mice but did not ameliorate the cachectic phenotype. In contrast, AB+G restored food intake and prevented loss of lean and fat mass. LLC AB+G mice exhibited reduced hypothalamic *Il6* and serotonin transporter (*Slc6a4*) expression, normalized adipocyte morphology and serum leptin levels, decreased adipose *Il6* and *Atgl* expression and reduced WAT sympathetic innervation. AB+G further lowered circulating corticosterone levels, and provided greater protection against muscle wasting, with increased *Pgc1α* expression and improved muscle function. Neither intervention affected tumor growth or tumor inflammatory gene expression.

**Conclusions:** Cancer cachexia is associated with a central and peripheral failure to appropriately activate AMPK signaling in response to the energetic stress imposed by cachexia. Combined AMPK activation and ghrelin administration exerted complementary effects on energy homeostasis, inflammation, and tissue wasting, resulting in greater protection against cachexia than either intervention alone. These findings support combined AMPK-ghrelin targeting as a promising therapeutic strategy for cancer cachexia.

## Introduction

Cancer-associated cachexia (CAC) is a multifactorial and systemic syndrome characterized by involuntary loss of skeletal muscle and adipose tissue depletion, which cannot be fully reversed by conventional nutritional interventions (1). This syndrome is associated with reduced survival, impaired quality of life, and poor clinical outcomes in patients with cancer (2). CAC is characterized by progressive weight loss, anorexia, fatigue, and profound metabolic disturbances, driven by a complex crosstalk between tumor-derived factors and host responses that promote inflammation and widespread metabolic dysregulation (1,2).

A hallmark of CAC is a persistent catabolic state in which protein and lipid breakdown exceed synthesis, leading to progressive wasting of skeletal muscle and white adipose tissue (3). In skeletal muscle, this involves enhanced proteolysis, reduced protein synthesis, and impaired regeneration, culminating in loss of mass and function (3,4). Concurrently, white adipose tissue undergoes increased lipolysis, impaired adipogenesis, and frequently increased thermogenesis (5,6). These catabolic processes are driven by tumor- and host-cachexokines (2,4).

Among inflammatory mediators, interleukin-6 (IL-6) plays a central role in CAC. Elevated circulating IL-6 levels correlate with the severity of weight loss, systemic inflammation, and poor prognosis in cancer patients (7,8). Experimental models demonstrate that increased levels of IL-6 in tumor-bearing mice drive muscle atrophy and adipose tissue remodeling by activating catabolic signaling and altering energy metabolism (6,9). Pharmacological or genetic inhibition of tumor-or host-derived IL-6 mitigates cachexia in preclinical studies, highlighting the IL-6/JAK/STAT3 pathway as a key link between tumor-driven inflammation and metabolic dysfunction (8–10).

Beyond peripheral effects, the central nervous system, particularly the hypothalamus, contributes to CAC by integrating peripheral inflammatory and metabolic signals to regulate appetite and energy balance (11,12). Tumor-induced neuroinflammation disrupts hypothalamic control, promoting anorexia, sympathetic activation, and altered energy metabolism (11,13). Cytokines such as IL-6 have been shown to alter hypothalamic appetite-regulating and neuroendocrine neuronal circuits, as well as metabolic sensing pathways (13,14).

In this context, AMP-activated protein kinase (AMPK) is of particular interest because it is a conserved energy sensor that coordinates cellular and systemic metabolism under energy stress (15). In the hypothalamus, AMPK modulates feeding behavior and energy expenditure particularly through neuropeptide regulation, while in peripheral tissues it regulates protein and lipid turnover (16,17). Although the role of AMPK in cancer-associated metabolic alterations remains poorly defined, and sometimes controversial, particularly in skeletal muscle (18–20), AMPK has emerged as a key regulator with anti-inflammatory effects (21,22), suggesting a protective role for AMPK activation in chronic inflammatory conditions such as CAC. Supporting this notion, skeletal muscle–specific deletion of AMPK accelerates age-related sarcopenia (23). Moreover, a recent study demonstrated that pharmacological stabilization of AMPK using a peptide-based approach prevents adipose tissue wasting in models of cancer cachexia (24). Yet, it remains unclear whether AMPK signaling is adequately engaged during CAC in response to the profound energy deficit associated with the syndrome. Furthermore, the role of AMPK in linking central and peripheral metabolic dysregulation with inflammation during cancer cachexia remains poorly understood.

Despite progress in deciphering CAC mechanisms, no pharmacological intervention has fully reversed the syndrome in humans (25,26). Here, we investigated hypothalamic and peripheral AMPK signaling during cancer cachexia and evaluated the therapeutic potential of pharmacological AMPK activation alone or in combination with the orexigenic/anabolic hormone ghrelin. Using the Lewis lung carcinoma (LLC), fibrosarcoma, and chronic viral infection-associated cachexia models (10,27), we showed that AMPK activation is blunted in both central and peripheral tissues across distinct models of established cachexia. Our results further demonstrated, in LLC-bearing mice, that pharmacological stimulation of AMPK, particularly when combined with ghrelin, attenuated tissue wasting, improved feeding behavior, and reduced circulating and tissue IL-6 signaling, supporting the therapeutic relevance of this combinatorial approach.

### Materials and methods Animals and Treatments

All procedures were approved by the Animal Care Ethics Committee of the University of Santiago de Compostela and followed European Directive 2010/63/EU. Male C57BL/6 mice (10–12 weeks of age) were housed under controlled environmental conditions (22 ± 2 °C, 12-hour light/dark cycle), with ad libitum access to chow and water.

To induce cachexia, 0.50 × 10⁶ Lewis lung carcinoma (LLC) cells suspended in 50 µL PBS were injected intramuscularly into the right thigh. Sham control mice received an equal volume of PBS. Once tumors became palpable (day 9), tumor-bearing mice were randomly assigned to the following experimental groups: (a) LLC, treated with vehicle only; (b) LLC AB, treated with AICAR (500 mg/kg, i.p.) and BC1618 (20 mg/kg, i.p.) administered on alternating days; (c) LLC AB+G, treated with AICAR, BC1618 and acylated ghrelin (0.8 mg/kg, s.c., twice daily);

LLC+G, treated with ghrelin alone (0.8 mg/kg, s.c., twice daily). Mice in the Sham group received the corresponding treatment vehicles.

Additional mouse models of cachexia, including fibrosarcoma CHX or MN/MCA1-bearing mice, and mice chronically infected with lymphocytic choriomeningitis virus (LCMV) were used to validate selected findings (see Supplementary Methods).

### Body Weight, composition, food intake and sample collection

Body weight was recorded every 2 days. Lean and fat mass were measured using EchoMRI before cancer cell implantation and at endpoint. Tumor mass was subtracted to calculate tumor-free body weight and lean mass. Food intake was assessed by manually weighing food pellets daily for 4 days.

Mice were euthanized 20 days after cancer cell implantation with isoflurane overdose followed by cervical dislocation. Blood was collected by retroorbital puncture, centrifuged (6,000 rpm, 10 min, 4 °C), and serum stored at −80 °C. Tissues collected included hypothalamus; gastrocnemius, anterior tibialis and soleus muscles from the non-tumor-bearing leg; gonadal and subcutaneous WAT, BAT, and tumor. Tissues were then either frozen or fixed in formaldehyde for subsequent molecular and histological analysis.

### Grip Strength

Forelimb grip strength was assessed using a grip strength meter. The results of five trials per mouse were averaged.

### Serum Analysis

Serum cytokines (IL-6, IL-1β, IFN-γ, TNF-α) and hormones (leptin, corticosterone) were measured in triplicate using ELISA or multiplex assays (Table S1).

### RNA Extraction and Quantitative PCR

Total RNA was extracted from frozen tissues using TRIzol reagent according to the manufacturer’s instructions. cDNA was synthesized using M-MLV reverse transcriptase. Quantitative real-time PCR was performed using Luminaris HiGreen Master Mix on a QuantStudio 7 Flex system. Gene expression was normalized to GAPDH (muscle/adipose) or HPRT (hypothalamus) and results were expressed as fold change to the control using the 2^−ΔΔCt method. Primer sequences are listed in Table S2.

### Western Blot

Proteins were extracted using lysis buffer for hypothalamic samples and RIPA buffer for muscle and adipose tissues. Samples were homogenized and centrifuged (13,000 rpm, 15 min, 4 °C). Protein concentration was determined using Bradford or BCA assays. Equal protein amounts were separated in SDS-PAGE gels, transferred to PVDF membranes, blocked with 5% BSA, and incubated overnight at 4 °C with primary antibodies (Table S3). HRP-conjugated secondary antibodies were applied, and signals detected using ECL. Total protein levels were normalized to β-actin (hypothalamus), GAPDH (muscle), or vinculin (WAT). Phosphorylated proteins were normalized to their respective total protein levels.

### Histology and Immunohistochemistry

Tissues were paraffin-embedded, sectioned (4 µm), and stained with hematoxylin and eosin according to standard histopathological techniques. Muscle fiber cross-sectional area (CSA) and adipocyte size were quantified from 3–4 random fields per sample using ImageJ, analyzing ≥25 fibers per field and 100 adipocytes per animal. For immunohistochemistry, sections underwent antigen retrieval, primary antibody incubation (e.g., anti-TH, 1:1000), HRP-conjugated secondary antibody, and DAB visualization. Images were acquired using a light microscope (Olympus IX73) and quantified using ImageJ software.

### iDISCO Clearing and Immunolabeling

Subcutaneous WAT was processed using the iDISCO+ method following the protocol developed by Renier et al (28). Samples were incubated with anti-UCP1 (1:1000) and anti-TH (1:2000) antibodies for 10 days at 37 °C, followed by incubation with Alexa Fluor–conjugated secondary antibodies for an additional 10 days. Cleared tissues were imaged using a Blaze light-sheet microscope (Miltenyi BioTec). Sympathetic innervation was quantified in five cubic regions per sample using Imaris software. Further details about procedure and quantification are shown in Supplementary methods.

### Statistical Analysis

Data are presented as mean and ±SEM. The number of animals included in each analysis is indicated as “*n*” in the corresponding figure legend. Statistical analyses were performed using GraphPad Prism 10 (GraphPad Software). Comparisons between two groups were conducted using an unpaired two-tailed Student’s *t*-test. Multiple groups were analyzed by one-way ANOVA, followed by Tukey’s test. Statistical significance was defined as p < 0.05.

## Results

### 1. LLC-bearing mice develop cachexia with upregulation of peripheral and hypothalamic IL-6

To characterize the metabolic and inflammatory alterations associated with cancer-associated cachexia (CAC), we performed a comprehensive phenotypic and molecular characterization of LLC-bearing mice at endpoint. Compared with shams, LLC mice exhibited significant reductions in body weight, lean mass, and fat mass (Fig. 1A), accompanied by reduced food intake (3.37 ± 0.10 vs 3.94 ± 0.07 g/day, p = 0.009, n = 9 mice per group). Serum analysis revealed elevated IL-6 and corticosterone levels, and reduced leptin concentrations, while TNF-α, IL-1β, and IFN-γ were unchanged (Table S4).

**Figure 1:**
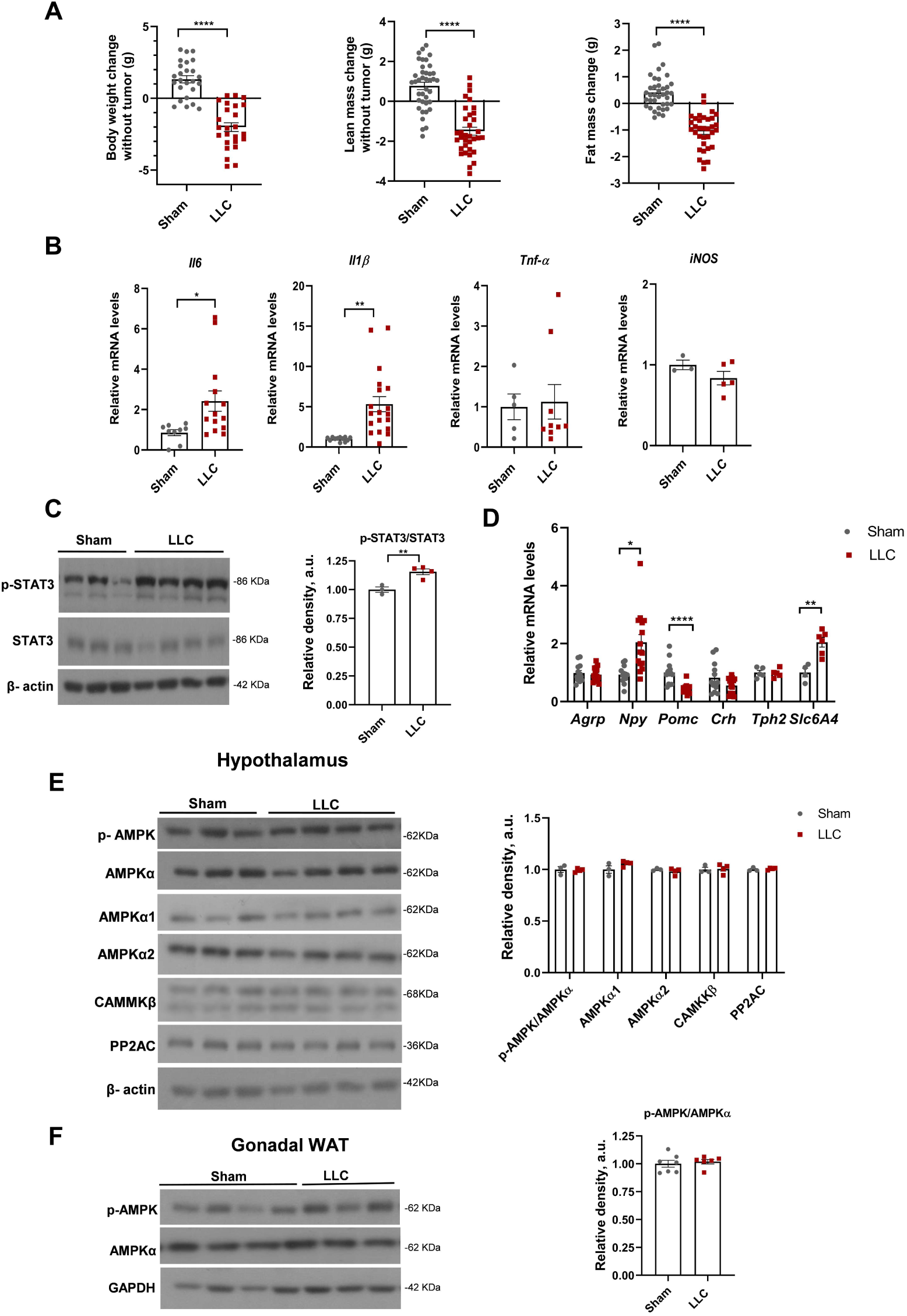
Lewis lung carcinoma (LLC)–bearing mice exhibit cachexia, hypothalamic inflammation, and impaired AMPK activation despite energy depletion. Mice were euthanized 20 days after the injection of cancer cells into the right hindlimb muscle (endpoint). Results for Sham and LLC mice (LLC) are shown. (**A**) Changes in body weight, lean mass, and fat mass. Final body weight and lean mass were calculated after subtraction of tumor mass. *n* = 34-39 mice per group. (**B**) Relative hypothalamic mRNA expression of *Il6*, *Il1β*, *Tnfα*, and *iNOS* assessed by RT-qPCR. *n* = 3-14 mice per group. (**C**) Hypothalamic pSTAT3 Tyr705 and total STAT3 protein levels analyzed by western blot. *n* = 3-4 mice per group (**D**) Relative hypothalamic mRNA levels of *Agrp, Npy, Pomc, Crh, Tph2* (Tryptophan Hydroxylase 2) and *Slc6a4* (serotonin transporter) quantified by RT-qPCR. *n* = 5-14 mice per group. (**E**) Protein levels of p-AMPKThr172, total AMPKα, AMPKα1 AMPKα2, CaMKKβ (calcium/calmodulin-dependent protein kinase kinase β) and PP2AC (Protein Phosphatase 2A Catalytic Subunit) in the hypothalamus and (F) in gonadal white adipose tissue (WAT) determined by western blot. *n* = 3-4 (E), 6-7 (**F**) mice per group Data are shown as mean ± SEM. Representative western blots are shown in (C), (E), and (F). Differences between groups were analyzed using an unpaired two-tailed Student’s *t* test. \**p* < 0.05, \*\**p* < 0.01, \*\*\**p* < 0.001, \*\*\*\**p* < 0.0001.

Skeletal muscle wasting was associated with upregulation of the atrophy-related genes *Murf1* and *Atrogin-1* and reduced expression of myosin heavy chain (*Mhc*). No significant changes were observed in the mRNA levels of *myogenin* or *Pgc1α* (Table S5).

In white adipose tissue (WAT), adipose triglyceride lipase (*Atgl*) mRNA was significantly increased, whereas hormone-sensitive lipase *(Hsl)* expression levels remained unchanged (Table S5), consistent with previous reports implicating ATGL as a key mediator of enhanced lipolysis in CAC (5,29). On the other hand, expression of browning-associated genes in WAT (*Ucp1, Cidea, Prdm16, Ppargc1α*) showed high interindividual variability and did not reach statistical significance (Table S5). Both skeletal muscle and WAT displayed increased *Il6* and *Il1β* expression (Table S5), suggesting local inflammatory responses.

In the hypothalamus *Il6* and *Il1β* mRNA levels were elevated, while *Tnf-α* and *iNOS* expression levels were similar to sham mice (Fig. 1B). Consistent with increased IL-6 signaling, phosphorylation of STAT3 was markedly elevated (Fig. 1C). Importantly, hypothalamic STAT3 activation was also observed in additional cachexia models, including fibrosarcoma (CHX)-bearing mice and mice with chronic lymphocytic choriomeningitis virus (LCMV) infection (Fig. S1A, B), indicating that activation of the IL-6/STAT3 pathway in the hypothalamus is a common feature of both tumor- and non-tumor-associated cachexia.

Because hypothalamic inflammation can disrupt energy-regulating circuits, we assessed the expression of key hypothalamic neuropeptides involved in feeding control (Fig. 1D). LLC-bearing mice showed increased mRNA levels of the orexigenic neuropeptide Y (*Npy)* and decreased levels of pro-opiomelanocortin (*Pomc*), the precursor of the anorexigenic neuropeptide α-MSH. Expression of agouti-related peptide (*Agrp*) and corticotropin-releasing hormone (*Crh*) remained unchanged. The reduced food intake, despite this pro-orexigenic transcriptional profile, is compatible with impaired secretion or signaling of these neuropeptides. We also examined the hypothalamic serotonergic pathway (Fig. 1D). LLC mice showed increased expression of the serotonin transporter *Slc6a4*, whereas the mRNA content of *Tph2*, the rate-limiting enzyme in serotonin synthesis, was unaffected. Increased *Slc6a4* expression may reflect enhanced serotonin reuptake, which has been associated with anhedonia, loss of appetite, and sickness behavior (30).

Together, these findings show that LLC-induced cachexia is characterized by loss of lean and fat mass, systemic inflammation, hypothalamic IL-6/STAT3 activation, dysregulation of hypothalamic neuropeptide and serotonergic signaling, as well as increased inflammatory gene expression in skeletal muscle and adipose tissue.

### 2. Impaired hypothalamic and peripheral AMPK activation in cachexia

AMPK is a central regulator of energy homeostasis that is normally activated during energy stress to restore ATP balance (15). In control mice, a 12-hour fast induced a robust increase in hypothalamic phosphorylation of AMPK at Thr172 (pAMPKThr172), confirming AMPK activation under conditions of low cellular energy (Fig. S2A). However, at endpoint, LLC-bearing mice displayed similar hypothalamic pAMPKThr172 levels to those of sham mice, despite exhibiting marked cachexia. (Fig. 1E). A similar impairment was observed in other cachexia models, including CHX-bearing mice and LCMV-infected mice (Fig. S2B, C), indicating that defective central AMPK activation is a common feature across distinct forms of cachexia.

To determine whether this defect resulted from changes in the content of AMPK or its regulatory machinery, we assessed the hypothalamic levels of AMPKα1 and AMPKα2 catalytic subunits, the upstream kinase CaMKKβ, and the phosphatase PP2AC. Levels of these proteins were comparable between LLC-bearing and sham mice (Fig. 1E), and similar results were observed in CHX and LCMV cachexia models (Fig. S2B,C). These data indicate that impaired hypothalamic AMPK activation is not attributable to reduced AMPKα protein abundance or changes in the levels of canonical regulators.

Likewise, in peripheral tissues, including WAT and liver, pAMPKThr172 levels did not differ between LLC and sham mice at endpoint (Fig. 1F and Fig. S3A), and the same pattern was obtained in CHX and MN/MCA1-bearing mice (Fig. S3C, D). Total AMPKα protein levels remained unchanged across all tissues analyzed (Figs. 1F, S3A-D).

These findings suggest that CAC is associated with a global failure of AMPK activation across central and peripheral tissues despite energy deficit.

### 3. AMPK activation preserves skeletal muscle but not adipose tissue in LLC mice

Given the critical roles of both central and peripheral AMPK in metabolic regulation, as well as its reported anti-inflammatory effects (21,22), we investigated whether pharmacological activation of AMPK could attenuate the cachectic phenotype. Two pharmacological activators with complementary mechanisms of action were selected: AICAR, a direct AMPK activator (31), and BC1618, a recently identified compound that inhibits proteasomal degradation of pAMPKThr172 (32).

When administered individually to LLC-bearing mice, neither AICAR (500 mg/kg, i.p.) nor BC1618 (20 mg/kg, i.p.) consistently increased hypothalamic AMPK phosphorylation or activated downstream targets such as ACC and Raptor (Fig. S4A, B). Moreover, treatment with either compound alone did not prevent the major cachexia-associated features of LLC-bearing mice (Fig. S4C).

To enhance AMPK phosphorylation, LLC mice were subsequently co-treated with AICAR and BC1618 administered on alternating days (AB treatment). This combined regimen significantly increased hypothalamic pAMPKThr172 and pRaptorSer792 in LLC mice at endpoint (Fig. 2A), indicating effective activation of central AMPK signaling. Increased pAMPKThr172 was also detected in skeletal muscle (Fig. 2B). In contrast, no increase in AMPK phosphorylation was observed in WAT of LLC AB mice (Fig. 2C), although AB treatment increased pAMPKThr172 levels in WAT of sham mice (Fig. S5A).

**Figure 2:**
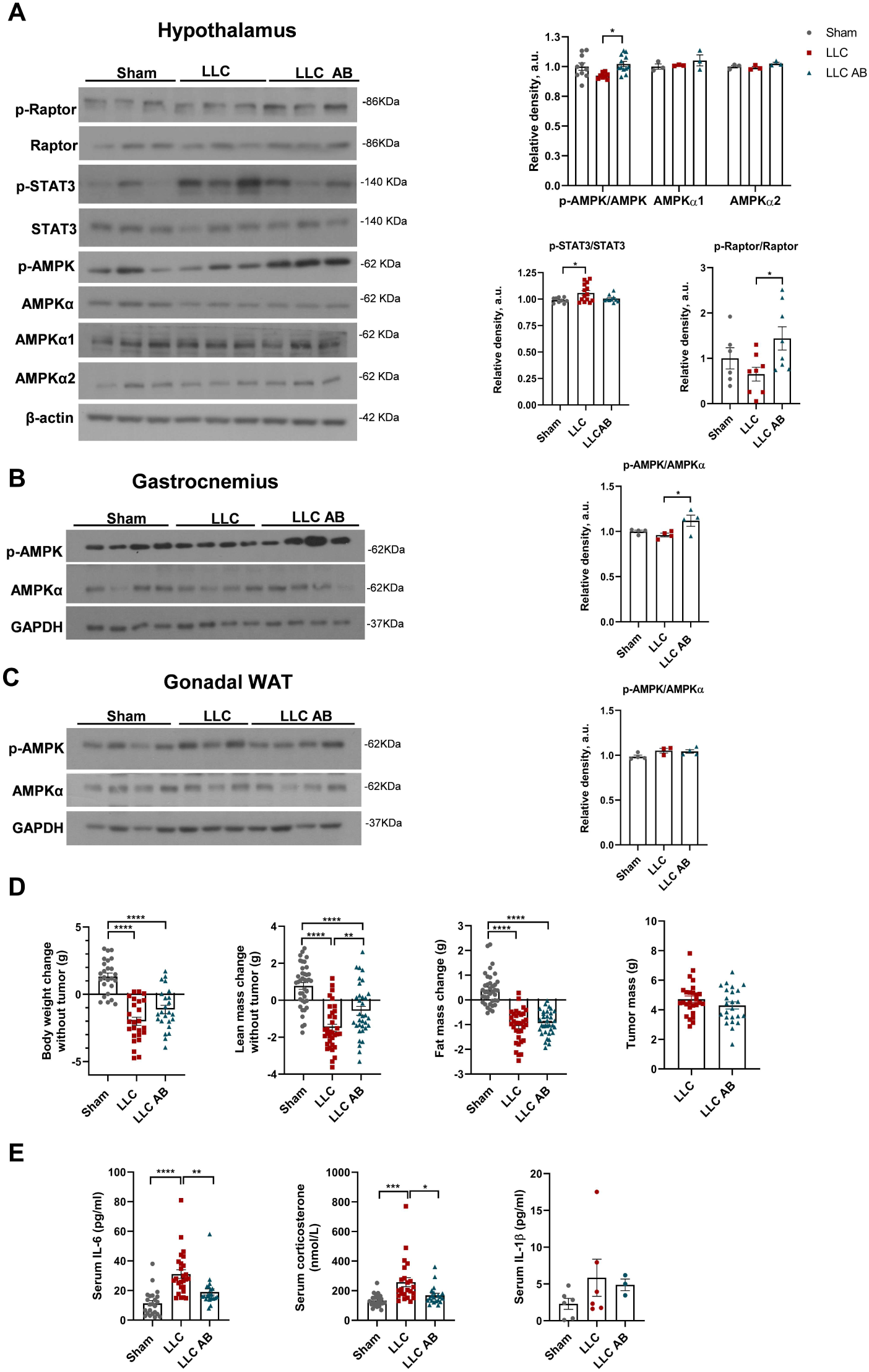
AICAR+BC1618 (AB) activates AMPK in the hypothalamus and skeletal muscle, but not adipose tissue in LLC-bearing mice. Results are shown for Sham, LLC, and AICAR+BC1618-treated LLC (LLC AB) mice at endpoint. Protein levels of pRaptorSer792, Raptor, pSTAT3Tyr705, STAT3, pAMPKThr172, total AMPKα, AMPKα1 and AMPKα2 assessed by western blot in the hypothalamus **(A)**, in the contralateral gastrocnemius muscle **(B)**, and in gonadal WAT **(C)**. *n* = 3-10 (A) and 3-4 (B, C) mice per group. **(D)** Changes in body weight, lean mass, and fat mass, as well as tumor mass. Final body weight and lean mass were calculated after subtraction of tumor weight. *n* = 34-39 mice per group. **(E)** Serum levels of IL-6, corticosterone, and IL-1β measured by ELISA. *n* = 3-23 mice per group. Data are presented as mean ± SEM. Representative western blots are shown in (A–C). Differences between groups were analyzed using one-way ANOVA, followed by Tukey’s test. \**p* < 0.05, \*\**p* < 0.01, \*\*\**p* < 0.001, \*\*\*\**p* < 0.0001.

Phenotypically, AB administration to LLC mice partially preserved lean mass but did not prevent the loss of body weight or fat mass (Fig. 2D). Notably, circulating IL-6 and corticosterone levels were significantly reduced compared with untreated LLC mice, whereas serum IL-1β levels remained high (Fig. 2E). Importantly, AB treatment did not affect LLC tumor mass (Fig. 2D) or tumor expression of *Il6* and *Il1β* (Fig. S5B), suggesting that its effects are mediated primarily through changes in host tissues rather than on the tumor itself.

Despite these systemic changes, AB-treated LLC mice remained hypophagic (Fig 3A). Likewise, hypothalamic neurotransmitter expression was similarly altered in treated and untreated LLC animals, with increased *Npy,* decreased *Pomc*, and elevated *Slc6a4* mRNA content (Fig. 3B). In addition, AB treatment did not significantly decrease hypothalamic *Il6* and *Il1β* mRNA levels (Fig. 3B).

**Figure 3:**
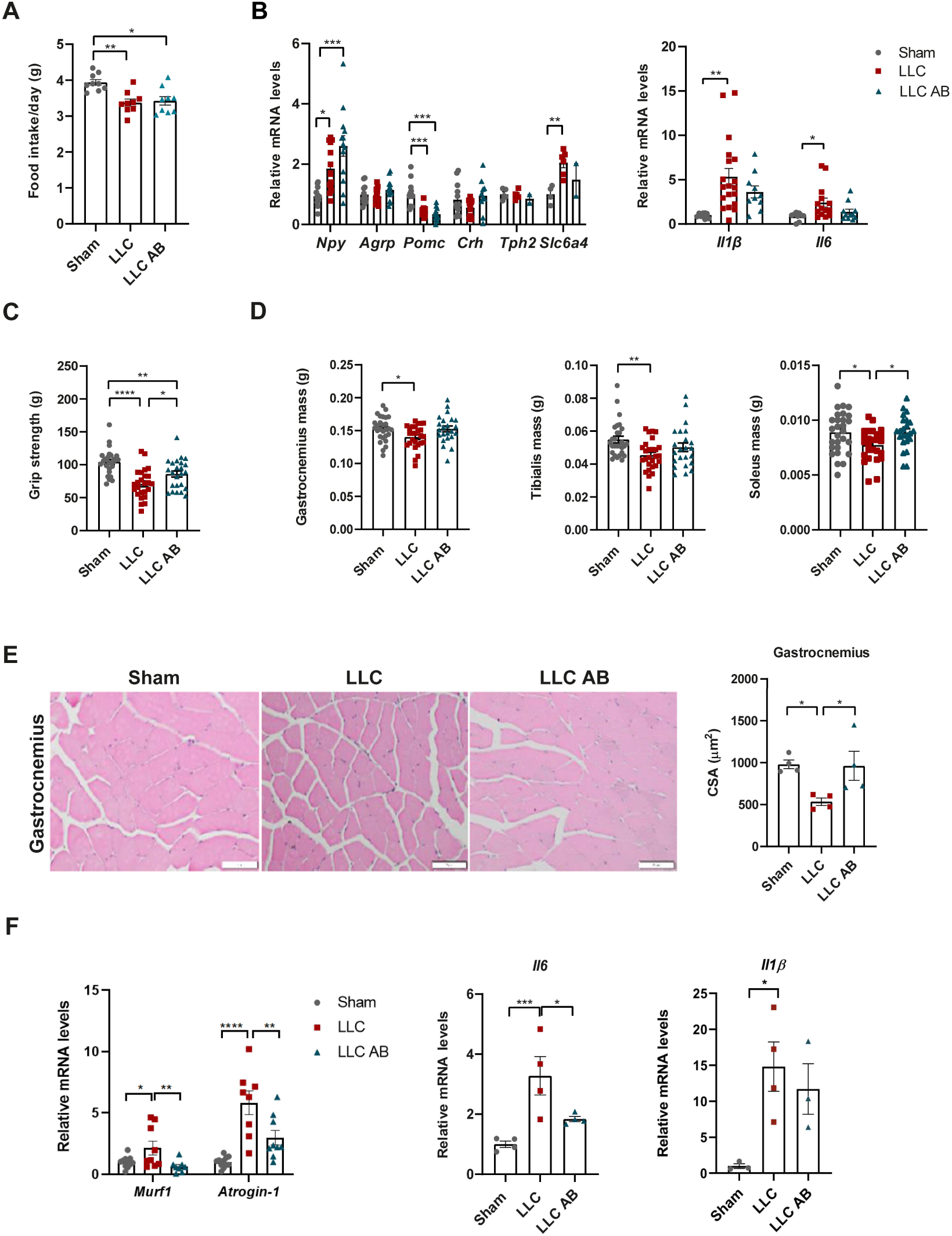
AICAR + BC1618 (AB) improves muscle function but fails to ameliorate hypophagia or hypothalamic inflammation. Results are shown for Sham, LLC, and LLC AB mice at endpoint. **(A)** Daily food intake. *n* = 9 mice per goup. **(B)** Relative hypothalamic mRNA levels of *Agrp, Npy, Pomc, Crh, Tph2* (tryptophan hydroxylase 2), *Slc6a4* (serotonin transporter), *Il6*, and *Il1β*, determined by qRT-PCR. *n* = 9-15 mice per group. **(C)** Grip strength. *n* = 24-26 mice per group. **(D)** Mass of contralateral gastrocnemius, anterior tibialis, and soleus muscles. *n* = 23-28 mice per group. **(E)** Cross-sectional area (CSA) of contralateral gastrocnemius muscle fibers quantified from hematoxylin and eosin–stained sections. *n* = 4 mice per group. **(F)** Relative mRNA expression of *Murf1, Atrogin-1*, *Il6* and *Il1β* in contralateral gastrocnemius muscle, determined by RT-qPCR. *n* = 3-5 mice per group. Data are presented as mean ± SEM. Differences between groups were analyzed using one-way ANOVA, followed by Tukey’s test. \**p* < 0.05, \*\**p* < 0.01, \*\*\**p* < 0.001, \*\*\*\**p* < 0.0001.

Regarding skeletal muscle function, LLC mice exhibited reduced forelimb grip strength, which was partially restored by AB treatment (Fig. 3C). This improvement was accompanied by preservation of skeletal muscle mass and fiber cross-sectional area (Fig. 3D, E and Fig. S5C). At the molecular level, AB reduced expression of *Murf1* and *Atrogin-1*, as well as *Il6* expression and STAT3 phosphorylation (Fig. 3F; Fig. S5D–F). Interestingly, muscle *Il1β* mRNA levels did not decrease (Fig. 3F), emphasizing that AB treatment primarily affects IL6 levels and signaling.

In contrast to skeletal muscle, adipose tissue loss persisted in AB-treated LLC mice. Thus, subcutaneous, and gonadal WAT, as well as brown adipose tissue mass, were still markedly reduced (Fig. 4A), and circulating leptin levels remained low (Fig. 4B). In addition, *Atgl* and *Hsl* expression, as well as ATGL protein levels, did not decrease in LLC AB compared with untreated LLC mice, while pHSL660 showed considerable variability among animals (Fig. 4C–E).

**Figure 4:**
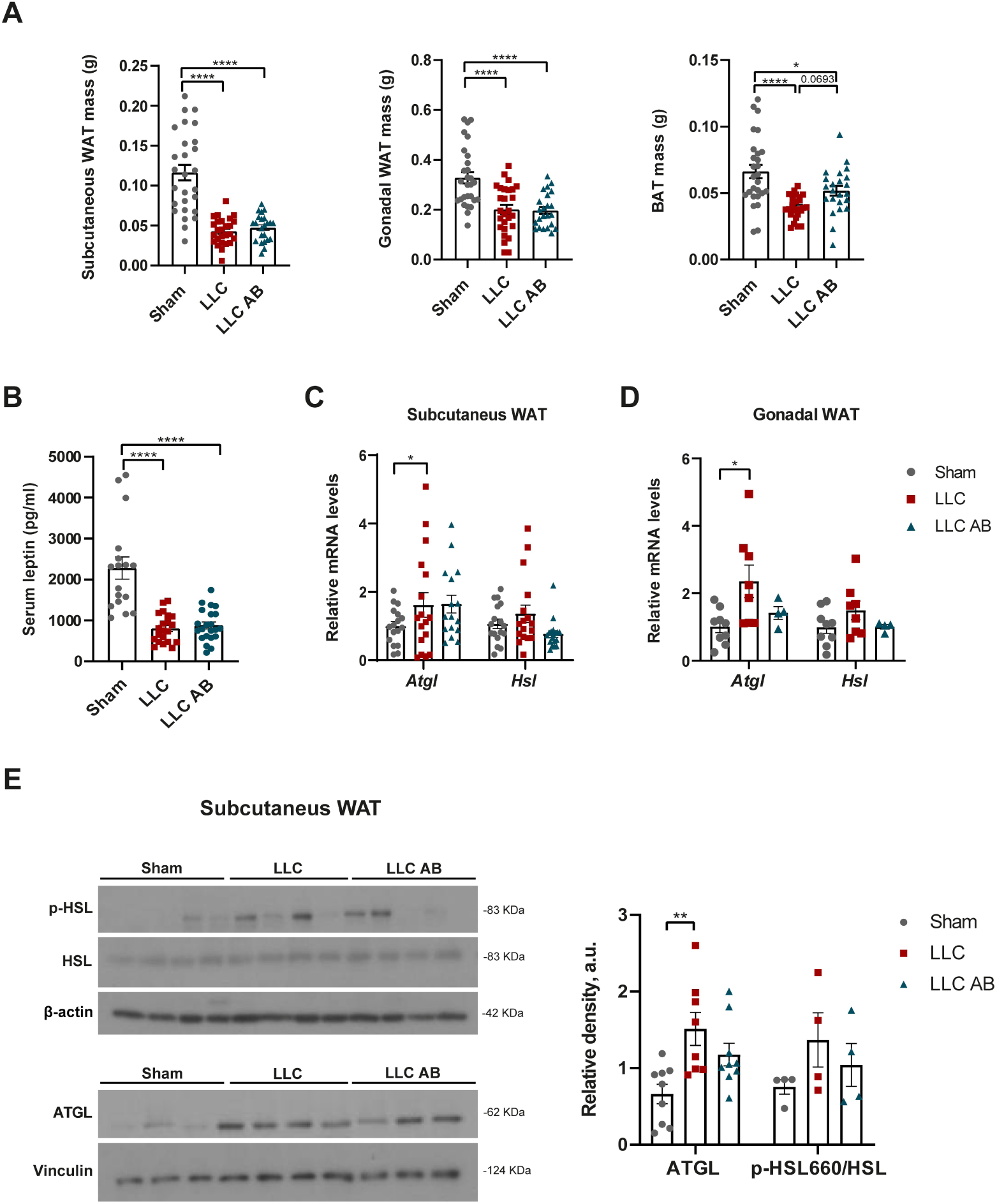
AICAR + BC1618 (AB) does not mitigate white adipose tissue (WAT) wasting in LLC-bearing mice. Results are shown for Sham, LLC, and LLC AB mice at endpoint. **(A)** Subcutaneous, gonadal, and brown adipose tissue mass. *n* = 21-28 mice per group. **(B)** Serum leptin levels determined by ELISA. *n* = 17-22 mice per group. **(C-D)** Relative mRNA levels of *Atgl* and *Hsl* in subcutaneous and gonadal WAT, determined by RT-qPCR. *n* = 16 (C) and 4-9 (D) mice per group. **(E)** Protein levels of ATGL, p-HSLSer660, and HSL in subcutaneous WAT, evaluated by western blot. Representative western blots are shown. *n* = 4-9 mice per group. Data are presented as mean ± SEM. Differences between groups were analyzed using one-way ANOVA, followed by Tukey’s test. \**p* < 0.05, \*\**p* < 0.01, \*\*\**p* < 0.001, \*\*\*\**p* < 0.0001.

Together, these results indicate that AB treatment in LLC mice activates AMPK in the hypothalamus and skeletal muscle and reduces systemic and skeletal muscle inflammation, as well as HPA axis activation, thereby preserving skeletal muscle mass and function. However, AB treatment does not restore food intake or prevent adipose tissue loss.

### 4. AMPK activation combined with ghrelin ameliorates cachexia

Given the limited effects of AMPK activation by AB alone on adipose tissue and feeding behavior in LLC mice, we next examined whether combining this treatment with the orexigenic/anabolic hormone ghrelin could further ameliorate the cachectic phenotype. This experiment included two additional groups: AB + ghrelin–treated LLC mice (LLC AB+G), and ghrelin-treated LLC mice (LLC+G) (Fig. 5A).

**Figure 5:**
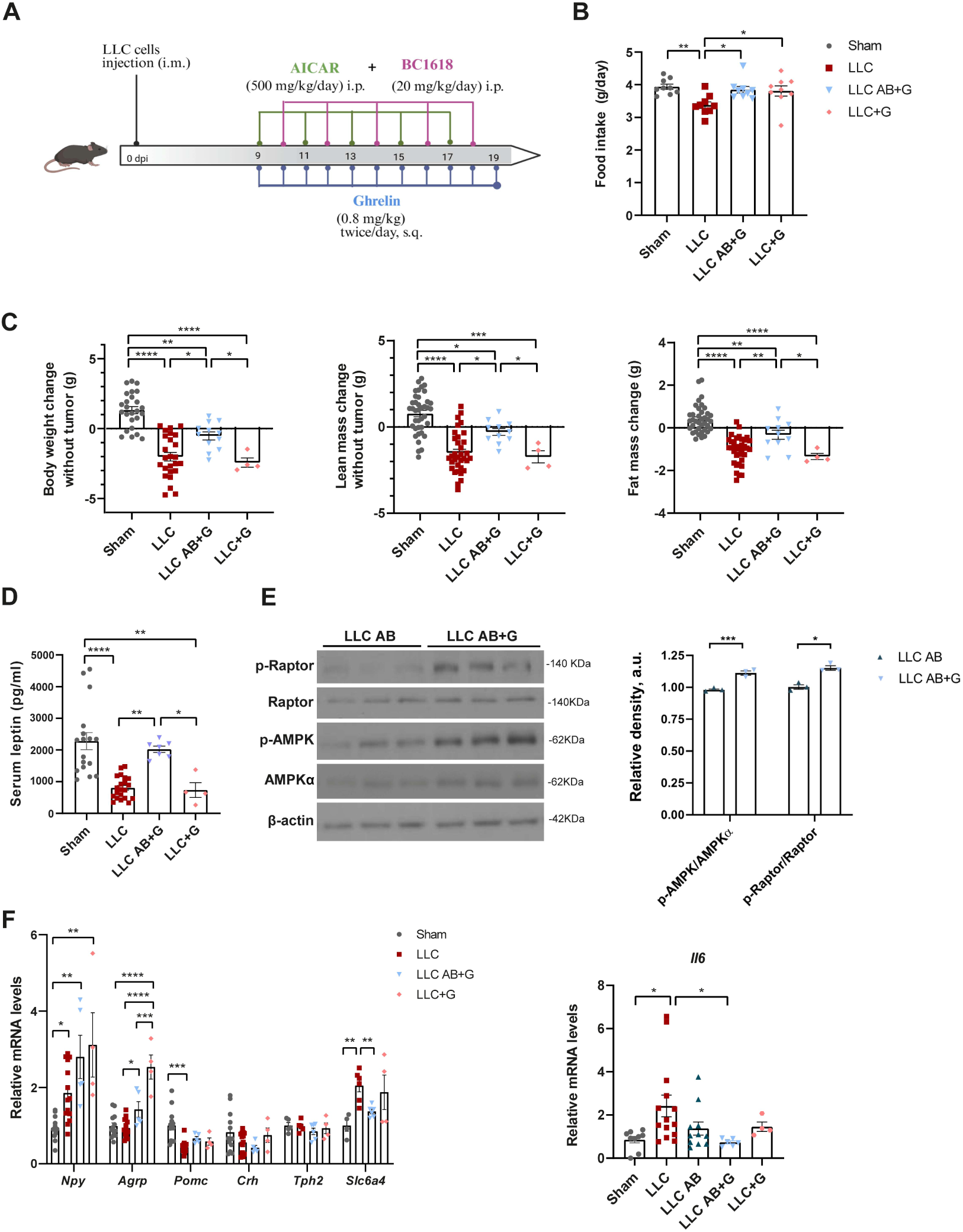
Combined AICAR + BC1618 and ghrelin (AB+G) treatment reduces body weight, muscle, and adiposity loss in LLC-bearing mice. Results are shown for Sham, LLC, LLC treated with AICAR + BC1618 + ghrelin (LLC AB+G), and LLC treated with ghrelin (LLC+G) mice at endpoint. **(A)** Schematic illustration of the treatment schedule with AB+G (Biorender). **(B)** Daily food intake. *n* = 9 mice per group. **(C)** Changes in body weight, lean mass, and fat mass. Final body weight and lean mass were calculated after subtracting tumor mass. *n* = 4-39 mice per group. **(D)** Serum leptin levels determined by ELISA. *n* = 4-17 mice per group. **(E)** Protein levels of p-AMPK Thr172, total AMPKα, Raptor and pRaptor Ser792 in the hypothalamus of LLC AB and LLC AB+G mice, evaluated by western blot. *n* = 3 mice per group. **(F)** Relative hypothalamic mRNA levels of *Npy, Agrp, Pomc, Crh, Tph2, Slc6a4* and *Il6* determined by RT-qPCR. *n* = 4-14 mice per group. Data are presented as mean ± SEM. Differences between groups were analyzed using one-way ANOVA, followed by Tukey’s test. \**p* < 0.05, \*\**p* < 0.01, \*\*\**p* < 0.001, \*\*\*\**p* < 0.0001.

LLC AB+G mice exhibited restored food intake to levels comparable to Sham controls (Fig. 5B), accompanied by recovery of body weight, lean mass, and fat mass relative to untreated LLC animals (Fig. 5C). Fat mass was also higher than in LLC AB animals (Table 1), and this effect was reflected by increased circulating leptin levels (Fig. 5D; Table 1). In contrast, LLC+G mice showed restoration of food intake, but no significant improvement in the other assessed parameters (Fig. 5B-D; Table 1).

**Table 1.** Differences between LLC AB, LLC AB+G and LLC+G mice.

|  | LLC AB | LLC AB+G | LLC+G |
| --- | --- | --- | --- |
| Body weight change (g) (n= 24 / 12 / 4) | -1.12 ± 0.30 | -0.60 ± 0.33 | -2.42 ± 0.32<sup>#</sup> |
| Lean mass change (g) (n= 34 / 11 / 4) | -0.57 ± 0.24 | -0.28 ± 0.20 | -1.73 ± 0.35<sup>#</sup> |
| Fat mass change (g) (n= 34 / 11 / 4) | -0.93 ± 0.08 | -0.32 ± 0.21<sup>**</sup> | -1.34 ± 0.14<sup>##</sup> |
| Food intake (g/day) (n= 9 / 9 / 9) | 3.42 ± 0.12 | 3.85 ± 0.09<sup>*</sup> | 3.80 ± 0.15 |
| Grip strength (g) (n= 24 / 12 / 4) | 85.98 ± 4.60 | 104 ± 4.40<sup>*</sup> | 83.89 ± 7.71<sup>#</sup> |
| Gastrocnemius mass (g) (n= 23 / 8 / 4) | 0.015 ± 0.004 | 0.016 ± 0.005 | 0.014 ± 0.005<sup>#</sup> |
| Tibialis mass (g) (n= 23 / 8 / 4) | 0.050 ± 0.002 | 0.058 ± 0.002 | 0.054 ± 0.006 |
| Soleus mass (g) (n= 23 / 8 / 4) | 0.008 ± 0.0003 | 0.009 ± 0.0008 | 0.008 ± 0.0005 |
| Gonadal WAT mass (g) (n= 23 / 8 / 4) | 0.19 ± 0.01 | 0.35 ± 0.02<sup>****</sup> | 0.19 ± 0.03<sup>##</sup> |
| Subc. WAT mass (g) (n= 23 / 8 / 4) | 0.047 ± 0.003 | 0.089 ± 0.008<sup>****</sup> | 0.040 ± 0.006<sup>###</sup> |
| BAT mass (g) (n= 23 / 9 / 4) | 0.05 ± 0.003 | 0.06 ± 0.003 | 0.04 ± 0.003<sup>#</sup> |
| Leptin (pg/ml) (n= 22 / 7 / 4) | 879 ± 83.57 | 2023 ± 99.8<sup>*</sup> | 734 ± 101<sup>#</sup> |
| Corticosterone (pg/ml) (n= 20 / 8) | 168.9 ± 14.11 | 82.80 ± 12.31<sup>**</sup> | N.D |
| IL-6 (pg/ml) (n= 22 / 9 / 4) | 19.02 ± 2.10 | 23.90 ± 1.55 | 31.73 ± 3.75<sup>*</sup> |
| IL-1β (pg/ml) (n= 3 / 5 / 3) | 4.88 ± 0.79 | 3.52 ± 1.31 | 3.26 ± 0.63 |

Parameters were assessed at the endpoint. Changes in body weight, lean mass, and fat mass were calculated as the difference (g) between endpoint and initial values for each animal. Data are presented as mean ± SEM. The number of animals per group (n) is indicated in parentheses for LLC AB, LLC AB+G and LLC+G mice. Statistical comparisons were performed using one-way ANOVA, followed by Tukey's post-hoc analysis. <sup>\*</sup>p<0.05, <sup>\*\*</sup>p<0.01, <sup>\*\*\*</sup>p<0.001, <sup>\*\*\*\*</sup>p<0.0001 versus LLC AB. <sup>#</sup>p<0.05, <sup>##</sup>p<0.01, <sup>###</sup>p<0.001 versus LLC AB+G.

At the hypothalamic level, AB+G-treated LLC mice exhibited greater AMPK activation than LLC AB animals (Fig. 5E). Interestingly, increased feeding in both ghrelin-treated LLC groups (LLC+G and LLC AB+G) was associated with elevated hypothalamic *Agrp* expression. However, only LLC AB+G mice normalized *Slc6a4* mRNa levels and reduced hypothalamic *Il6* mRNA content (Fig. 5F).

AB+G also restored subcutaneous and gonadal fat depots in LLC mice (Fig. 6A, B; Table 1) and histological analyses revealed normalization of adipocyte morphology (Fig. 6C). Molecular studies showed reduced ATGL expression with no change in HSL mRNA (Fig. 7A). Additionally, AB+G reduced *Il6* expression, which may contribute to the increased pAMPK levels observed in fat (Fig. 7C). Note that AB alone did not decrease WAT *Il6* mRNA, and neither AB+G nor AB was able to attenuate *Il1β* expression (Fig. 7B). These findings support an interaction between AB and ghrelin in restoring adipose tissue metabolism and inflammation in LLC mice.

**Figure 6:**
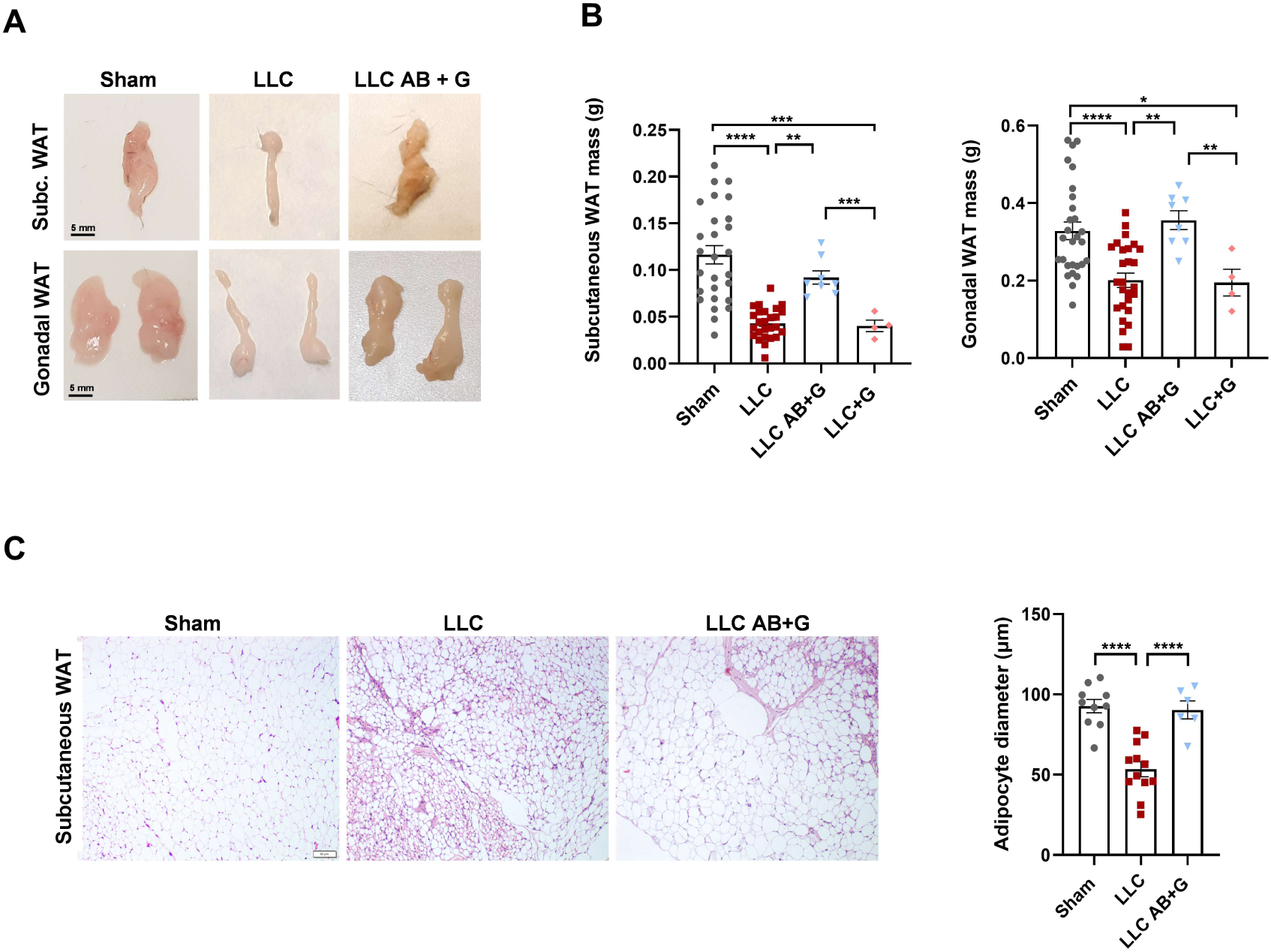
Treatment with AICAR + BC1618 and ghrelin (AB+G) prevents subcutaneous and gonadal WAT mass loss in LLC-bearing mice. Results for Sham, LLC, LLC AB+G, and LLC+G mice at endpoint are shown. **(A)** Representative photographs of subcutaneous (top) and gonadal (bottom) adipose tissue. **(B)** Subcutaneous and gonadal WAT mass. *n* = 4-28 mice per group. **(C)** Adipocyte diameter in subcutaneous WAT quantified from hematoxylin and eosin–stained tissue sections. *n* = 6-12 mice per group. Data are presented as mean ± SEM. Differences between groups were analyzed using one-way ANOVA, followed by Tukey’s test. \**p* < 0.05, \*\**p* < 0.01, \*\*\**p* < 0.001, \*\*\*\**p* < 0.0001.

**Figure 7:**
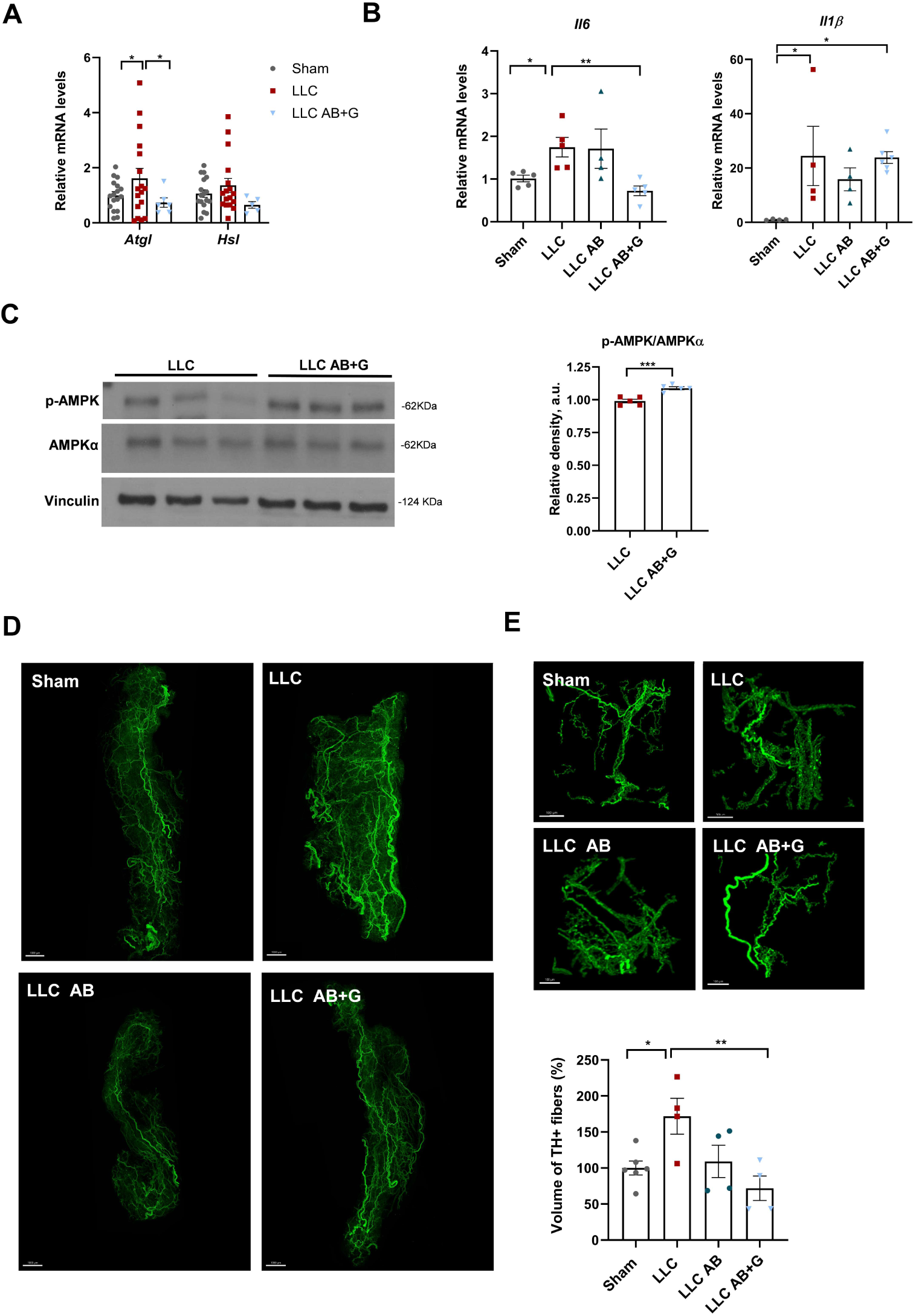
AICAR + BC1618 + ghrelin (AB+G) in LLC-bearing mice reduces ATGL expression, IL-6 levels, and sympathetic innervation in WAT. Results in subcutaneous WAT from Sham, LLC, and AICAR+ BC1618+ ghrelin-treated LLC mice (LLC AB+G) mice at endpoint are shown. **(A)** Relative mRNA levels of ATGL and HSL determined by RT-qPCR. *n* = 5-17 mice per group. **(B)** Relative expression of *Il6* and *Il1β* assessed by RT-qPCR. *n* = 4-6 mice per group **(C)** Protein levels of pAMPKThr172 and total AMPKα levels from LLC and LLC AB+G mice evaluated by western blot. A representative western blot is shown. *n* = 5 mice per group. **(D)** TH-positive innervation in whole subcutaneous WAT, visualized after iDISCO clearing, and quantification of the percentage volume of TH-stained fibers using Imaris. **(E)** For quantification, the volume of TH-positive fibers was determined in 5 randomly selected tissue cubes from the subcutaneous WAT of each mouse. A representative 3D-reconstruction of TH positive fibers of one cube per animal group is shown. *n* = 4-6 mice per group. Data are presented as mean ± SEM. Differences between groups were analyzed using one-way ANOVA, followed by Tukey’s test. \**p* < 0.05, \*\**p* < 0.01, \*\*\**p* < 0.001, \*\*\*\**p* < 0.0001.

Because sympathetic nervous system activity regulates adipose tissue lipolysis, we examined sympathetic innervation of WAT by quantifying tyrosine hydroxylase (TH)-positive fibers as a marker of noradrenergic nerve density. Both, tissue-section tyrosine hydroxylase (TH) immunohistochemistry (Fig. S6A) and whole-tissue clearing by iDISCO combined with light-sheet microscopy (Fig. 7D, E) revealed increased TH–positive innervation in subcutaneous WAT of LLC mice. This increase was significantly reduced by AB+G, but not by AB treatment (Fig. 7D, E, Fig. S6A, B, movie S1).

Finally, AB+G administration in LLC mice provided greater protection against skeletal muscle wasting than AB alone. Muscle mass, fiber cross-sectional area, and grip strength were markedly improved (Fig. 8A-C; Table 1) accompanied by reduced *Murf1, Atrogin-*1, and *Il6* expression, whereas *Il1β* mRNA levels remained unaltered (Fig. 8D). AB+G also increased *Pgc1α* expression (Fig. 8D), suggesting activation of a PGC-1α–dependent program promoting mitochondrial biogenesis and suppression of muscle atrophy pathways. Furthermore, although circulating IL-6 remained low and comparable to those in LLC AB mice (Table 1), corticosterone levels were further reduced (Fig. 8E; Table 1). Notably, tumor mass and tumor *Il6* and *Il1b* expression were unaffected (Fig. 8F).

**Figure 8:**
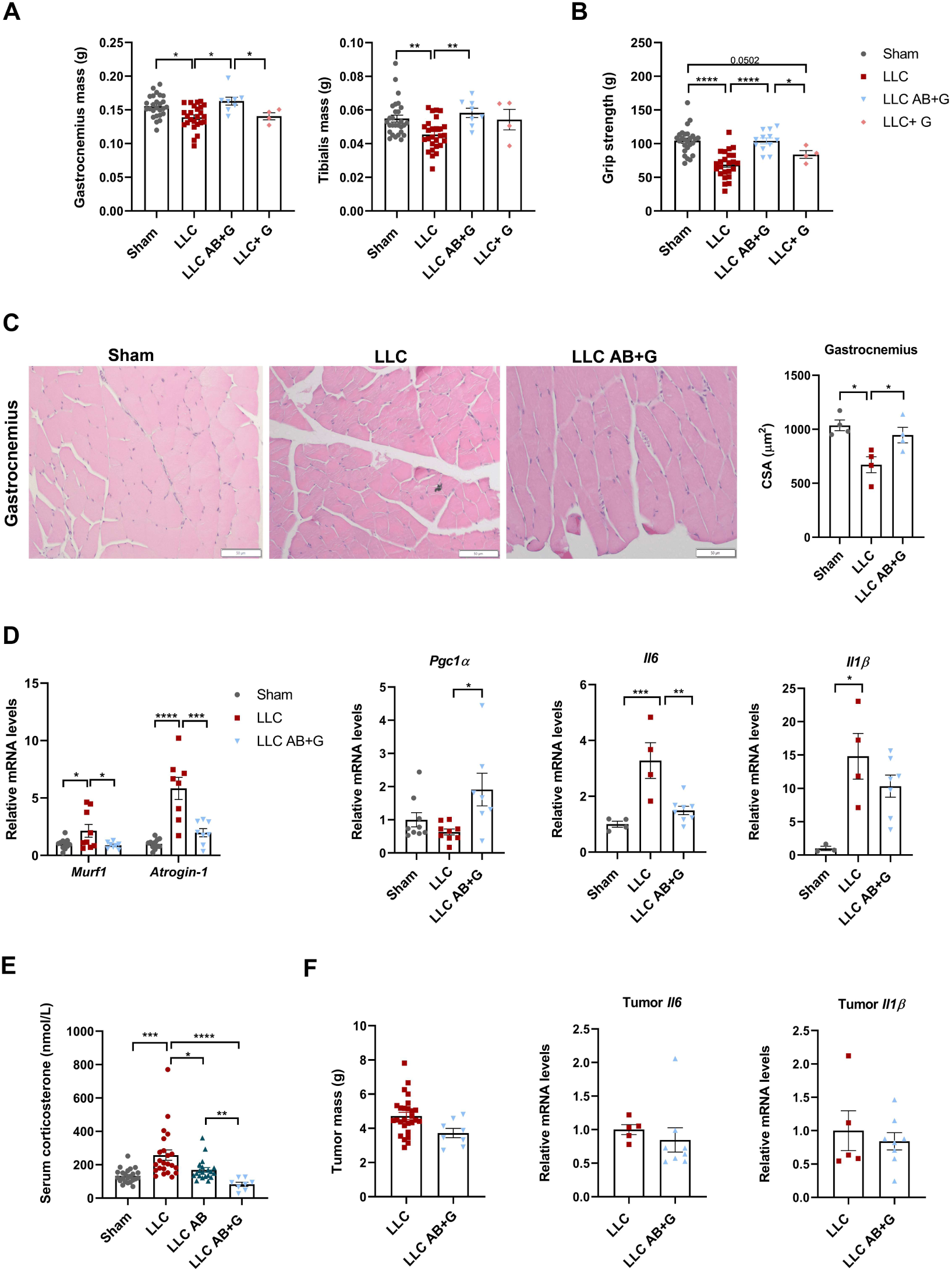
AICAR + BC1618 + ghrelin (AB+G) markedly improves muscle mass and function in LLC-bearing mice. Results for Sham, LLC, AICAR + BC1618 + ghrelin-treated LLC (LLC AB+G), and LLC-treated with ghrelin (LLC+G) mice at endpoint are shown. **(A)** Contralateral gastrocnemius and tibialis mass. *n* = 4-28 mice per group. **(B)** Grip strength. *n*= 4-28 mice per group. **(C)** Contralateral gastrocnemius muscle fiber cross-sectional area (CSA) quantified from hematoxylin and eosin–stained histological sections. *n* = 4 mice per group. **(D)** Relative mRNA levels of *Murf1*, *atrogin1*, *Pgc1α, Il6* and *Il1β* determined by RT-qPCR. *n* = 4-11 mice per group. **(E)** Serum corticosterone levels assessed by ELISA. *n* = 8-23 mice per group. **(F)** Tumor mass, and relative mRNA levels of *Il6* and *Il1β* in tumors from LLC and LLC AB+G mice determined by RT-qPCR. *N* = 5-20 mice per group. Data are presented as mean ± SEM. Differences between groups were analyzed using one-way ANOVA, followed by Tukey’s test. \**p* < 0.05, \*\**p* < 0.01, \*\*\**p* < 0.001, \*\*\*\**p* < 0.0001.

Collectively, these findings identify impaired central and peripheral AMPK signaling in CAC and support combined AMPK and ghrelin targeting as an effective therapeutic strategy.

## Discussion

Cancer cachexia is a complex metabolic syndrome characterized by involuntary loss of skeletal muscle and adipose tissue driven by systemic inflammation and profound dysregulation of energy homeostasis (1,2). Despite its clinical relevance, effective therapies remain limited, and current interventions are largely supportive rather than disease-modifying.

In this study, we used the Lewis lung carcinoma (LLC) model in C57BL/6 mice, together with additional cachexia models, to investigate the role of energy-sensing and neuroendocrine pathways. Consistent with previous reports, LLC-bearing mice developed marked loss of lean and fat mass accompanied by systemic and tissue inflammation, including elevated circulating IL-6 and IL-1β and increased local inflammatory gene expression across key metabolic tissues.

LLC mice exhibited reduced food intake, although hypothalamic neuropeptide expression indicated a compensatory orexigenic response, with increased *Npy* and reduced *Pomc* expression. This dissociation between neuropeptide transcription and feeding behavior potentially reflects impaired neuropeptide release or signaling. The increased hypothalamic serotonergic reuptake may further contribute to anorexia and sickness behavior (30). Elevated hypothalamic expression of *Il1β* and *Il6* is likely related to these alterations and to the increased levels of circulating corticosterone in CAC mice (11,13,33). Together, these data support a coordinated inflammatory and neuroendocrine response contributing to anorexia and metabolic dysfunction in CAC.

A central finding of this study is a failure to appropriately engage AMPK signaling in response to the severe energetic stress associated with cachexia. Under physiological conditions, AMPK is activated during energy deprivation to restore metabolic homeostasis (15). In contrast, AMPK phosphorylation does not increase in hypothalamus, skeletal muscle, and peripheral metabolic tissues across multiple cachexia models (LLC, CHX, and LCMV), despite profound systemic energy imbalance.

Importantly, in contrast to previous reports describing reduced AMPKα protein levels in adipose tissue during cancer cachexia (24), we did not observe changes in AMPKα abundance, supporting the concept that AMPK dysregulation in cachexia may also reflect impaired signaling activation or regulation, in a tissue-and context-dependent manner. We did not find decreased levels of canonical regulatory components, including CaMKKβ or PP2AC. One potential explanation is that inflammatory and stress-related pathways may interfere with AMPK activation or downstream signaling, thereby limiting its adaptive response under cachectic conditions. Alternative, enhanced turnover of phosphorylated AMPK may contribute, since Thr172 phosphorylation can promote recognition by ubiquitin-dependent pathways such as Fbxo48-mediated proteasomal degradation (32), although this requires further validation in cachexia models.

Collectively, these data support the concept that cancer cachexia is associated with a maladaptive energy-sensing response.

Given the anti-inflammatory role of AMPK, including inhibition of NF-κB and JAK/STAT signaling pathways (21,22,34), we next investigated whether pharmacological AMPK activation could attenuate cachexia. We used AICAR, a direct AMPK activator (31), and BC1618, which prolongs AMPK activation by preventing degradation of phosphorylated AMPK (32). Administration of either compound alone resulted in only modest effects and did not significantly improve muscle or fat loss in LLC-bearing mice. This contrasts with previous findings in C26 and other cachexia models, where AICAR ameliorated body weight and lean mass (19), suggesting experimental differences.

To achieve stronger AMPK activation, we combined AICAR and BC1618 (AB). This regimen activated AMPK in the hypothalamus and skeletal muscle, but not in white adipose tissue, indicating tissue-specific differences in AMPK responsiveness during cachexia. Functionally, AB treatment attenuated skeletal muscle wasting and improved muscle performance but did not restore food intake or prevent adipose tissue loss. Tumor growth was unaffected, indicating that the observed effects were mediated by host metabolic modulation rather than anti-tumor activity.

Mechanistically, AB treatment reduced circulating IL-6 and corticosterone levels and suppressed *Il6* expression and STAT3 activation in skeletal muscle. These findings suggest that attenuation of the IL-6/STAT3 pathway contributes to the protective effects of AB in skeletal muscle. Potential mechanisms include reduced *Il6* transcription and inhibitory phosphorylation of JAK1 by AMPK, thereby limiting STAT3 activation (35,36). Overall, these anti-inflammatory effects likely outweigh the inhibitory effects of AMPK on protein synthesis described in non-cachectic settings (20). These results support a predominant muscle-specific effect of AB induced-AMPK activation in attenuating skeletal muscle wasting in CAC, although minor neuroendocrine contributions cannot be excluded, such as the reduction of corticosterone levels

Despite these beneficial effects, AB treatment did not activate AMPK in WAT, and inflammatory signaling remained elevated, as indicated by persistent *Il6* expression and STAT3 activation. Consistently, fat loss continued and was associated with increased *Atgl* expression. These data suggest that adipose tissue in CAC is more resistant to pharmacological AMPK activation.

With the aim of preserving muscle mass while restoring appetite and fat mass, we next combined AB treatment with ghrelin (AB+G). Ghrelin has well-established orexigenic, anabolic, and anti-inflammatory properties (37). Several studies have reported beneficial effects in experimental cachexia (29,37), although clinical evidence remains inconsistent, and the ghrelin receptor agonist anamorelin has not been approved by the FDA or EMA for CAC treatment, although it has been authorized in Japan for certain tumor-induced cachexia (38).

In our study, ghrelin alone increased food intake but did not significantly improve the cachectic phenotype of LLC mice. In contrast, AB+G markedly ameliorated cachexia. LLC AB+G mice showed reduced body weight loss and restoration of both lean and fat mass, together with normalization of food intake. Importantly, the inability of ghrelin alone to reproduce these effects despite increasing food intake suggests that the benefits of AB+G cannot be explained solely by enhanced caloric intake and likely involve complementary metabolic, neuroendocrine, and anti-inflammatory mechanisms. The beneficial effects of AB+G were associated with enhanced hypothalamic AMPK activation, increased *Agrp* expression, reduced hypothalamic *Il6* levels, and normalization of serotonergic signaling, suggesting improved central regulation of energy balance and inflammation.

Importantly, the combined therapy also restored adipose tissue mass, an effect not observed with either treatment alone. This was associated with increased AMPK activation, reduced *Atgl* and *Il6* expression and decreased STAT3 activation in WAT. These findings suggest that concurrent central and local modulation of inflammation may be important for preserving adipose tissue in CAC.

In WAT of LLC mice we also observed increased sympathetic innervation, consistent with previous reports (6,39). Given the role of the sympathetic nervous system in regulating lipolysis and thermogenesis, this likely contributes to fat loss in cachexia. Using whole-tissue clearing and light-sheet microscopy, we found that AB+G treatment, effectively reduced sympathetic fiber density in subcutaneous adipose tissue of LLC mice, suggesting that AB+G partially reversed cachexia-associated sympathetic remodeling. This effect may result from reduced local inflammation and modulation of hypothalamic circuits controlling sympathetic outflow.

Finally, the combined therapy reinforced skeletal muscle protection. It preserved muscle mass and fiber size, improved muscle function, suppressed atrogenes and increased *Pgc1α* expression, indicating improved metabolic and mitochondrial function. These changes were accompanied by further reductions in circulating corticosterone, suggesting improved systemic neuroendocrine homeostasis.

Several limitations should be acknowledged. Although pharmacological modulation of AMPK signaling attenuated multiple cachectic features and was associated with beneficial metabolic and anti-inflammatory effects, the present study does not establish a definitive causal role for AMPK in mediating these responses. Future tissue-specific genetic studies will be required to define the contribution of central and peripheral AMPK signaling to these effects.

In conclusion, our findings demonstrate that cancer cachexia is associated with impaired AMPK responsiveness in both central and peripheral tissues despite severe energetic stress. Pharmacological AMPK activation partially attenuates cachexia by reducing inflammation and muscle catabolism but is insufficient to prevent adipose tissue loss or restore appetite. Importantly, combining AMPK activation with ghrelin exerts complementary effects on energy homeostasis, inflammation, and tissue wasting, resulting in greater protection against cachexia than either intervention alone. These results support a multimodal therapeutic strategy targeting both energy sensing and neuroendocrine regulation as a promising approach for cancer-associated cachexia.

## Acknowledgments

The authors gratefully acknowledge the excellent technical assistance of Luz Casas and Juan Silva (Dep. Physiology. University of Santiago de Compostela), and Dr. Jose Luís Pardo-Vázquez (University of A Coruña). We also thank Prof. R. Zechner and Prof. A. Bergthaler for generously providing samples from cachectic mice used in this study. The authors of this manuscript certify that they comply with the ethical guidelines for authorship and publishing in the Journal of Cachexia, Sarcopenia and Muscle (40).

## Author contributions

**R.S.:** conceptualization. **V.G-A., S.C., A.R., J.C-M., A.C-D., N.S., A.P-L., P.B., M.S.:** performed research. **D.V., A.V., F.TA., VM. A, and R.S.:** supervision and analysis of data. **R.S. and V.G-A:** wrote original draft. **V.G-A., A.R., D.V., A.V., F.T-A., VM. A., and R.S.:** reviewed and edited manuscript. **R.S.:** funding acquisition and project administration. All authors contributed to the article and approved the submitted version

---

**Suppl. Fig. 1:**
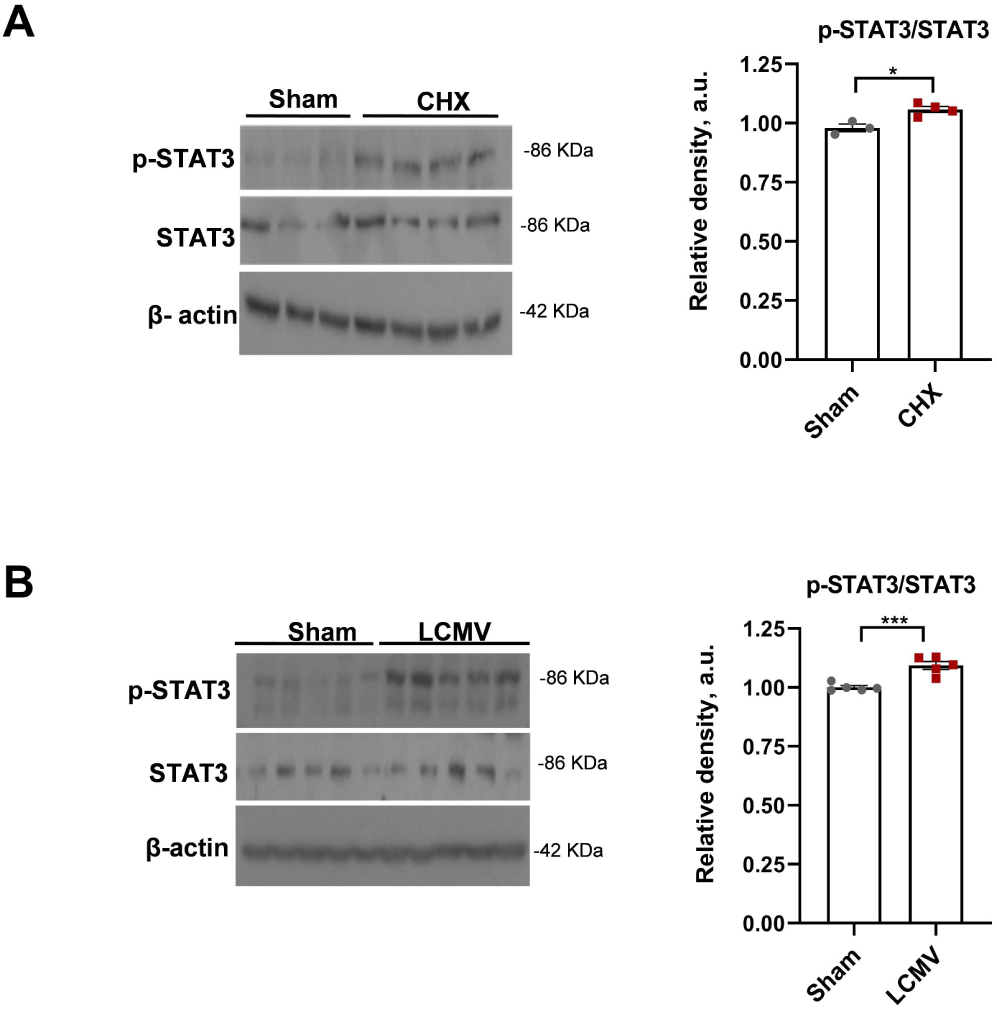
**(A, B)** Levels of pSTAT3Tyr705 and STAT3 in the hypothalamus of CHX fibrosarcoma–bearing mice and Lymphocytic choriomeningitis virus (LCMV)-infected mice, respectively, assessed by Western blot. *n* = 3–4 (A), 5 (B) mice per group. Data are presented as mean ± SEM. Differences between groups were analyzed using an unpaired two-tailed Student’s *t* test. \**p* < 0.05, \*\*\**p* < 0.001

**Suppl. Fig. 2:**
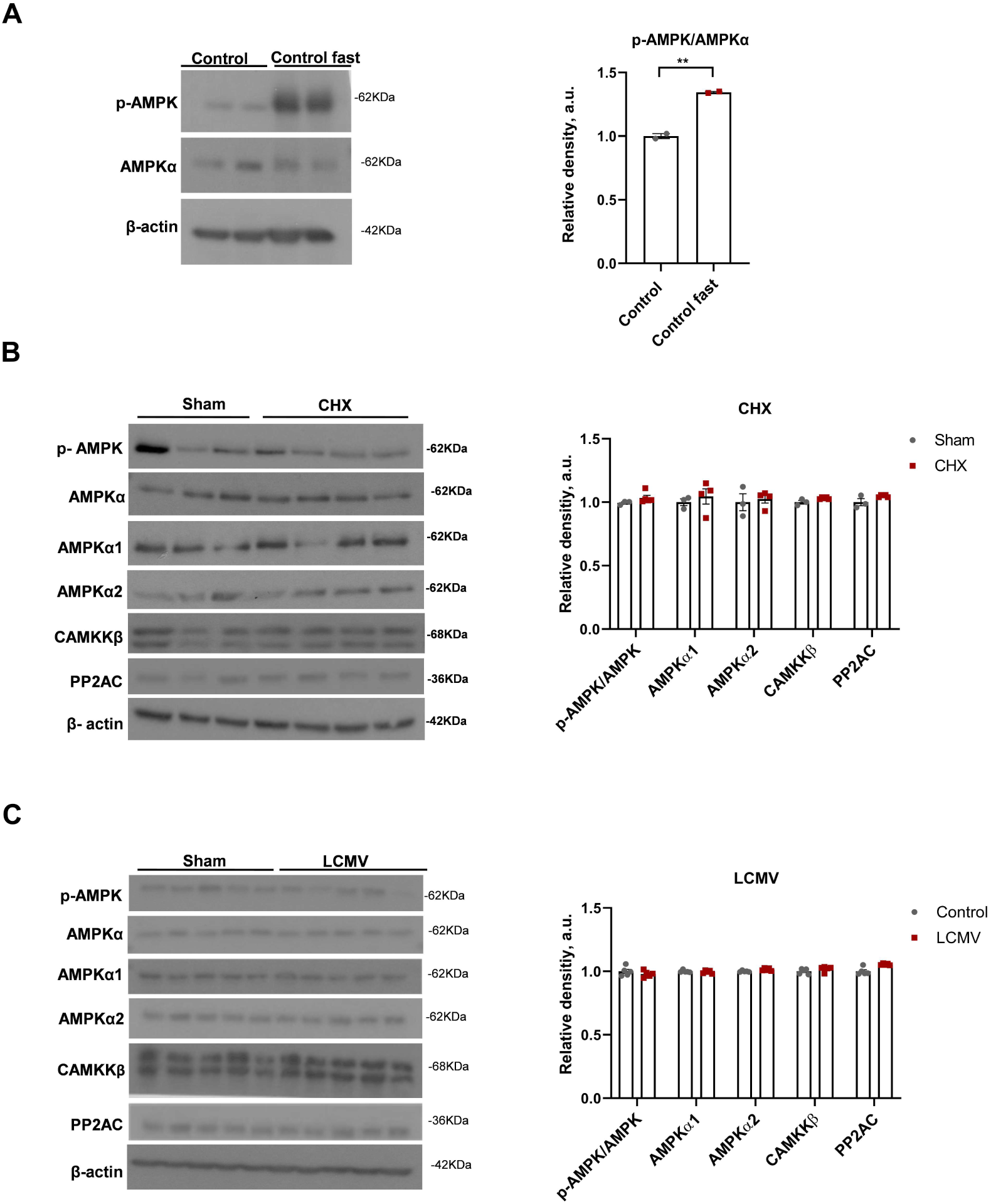
**(A)** Levels of pAMPKThr172 and total AMPKα in control mice and mice fasted overnight determined by western blot. **(B, C)** Levels of p-AMPKThr172, total AMPKα, AMPKα1, AMPKα2, CaMKKβ (calcium/calmodulin-dependent protein kinase kinase β), PP2AC (Protein Phosphatase 2A catalytic subunit) in the hypothalamus of CHX fibrosarcoma-bearing mice and mice infected with Lymphocytic choriomeningitis virus (LCMV), respectively, studied by western blot. Data are presented as mean ± SEM. *n* = 3 mice per group in (A), *n*= 3-4 in (B) and *n*=5 in (C). Differences between groups were analyzed using an unpaired two-tailed Student’s *t* test. \*\**p* < 0.01.

**Suppl. Fig. 3:**
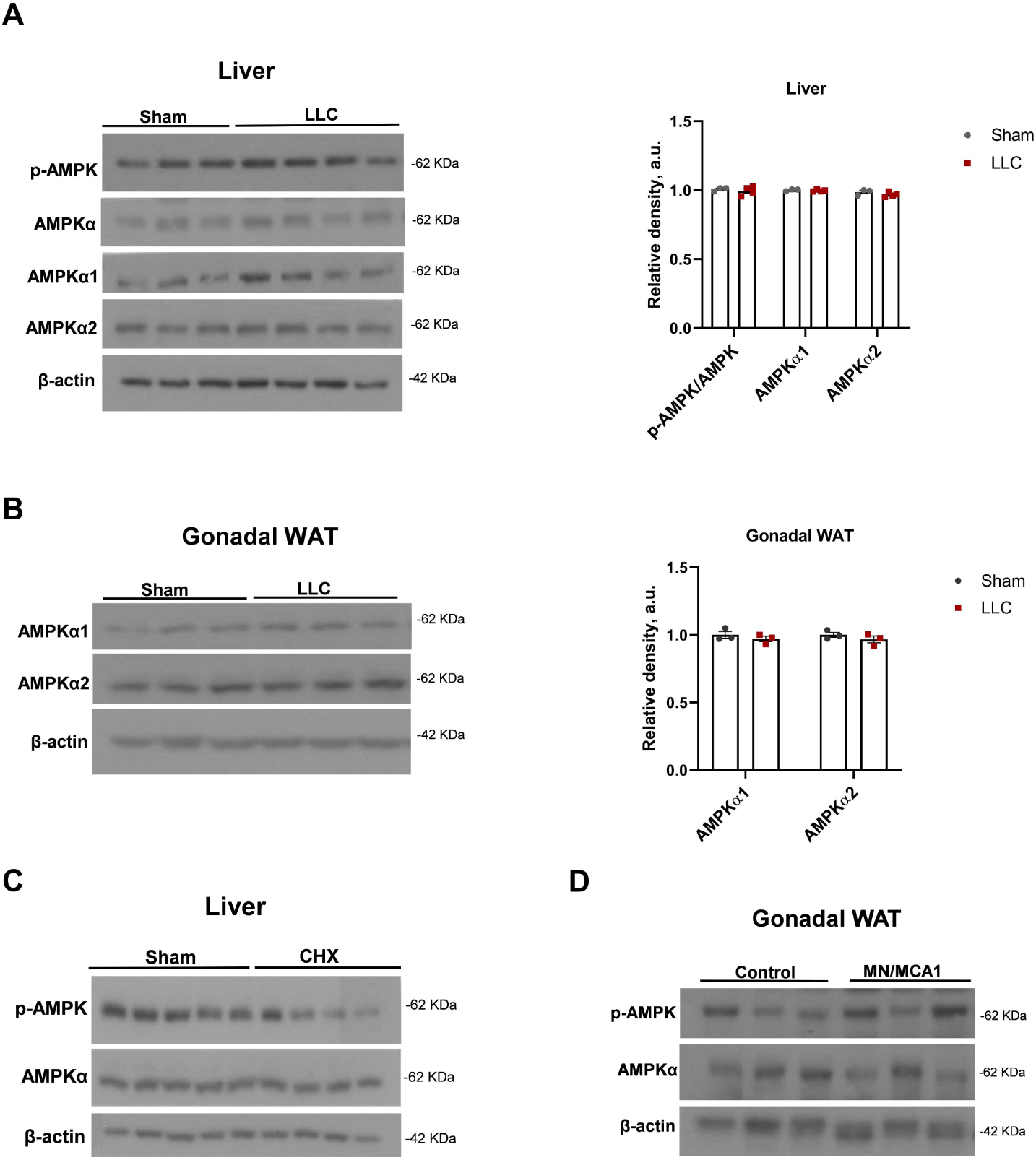
Levels of pAMPKThr172, total AMPKα, AMPKα1 and AMPKα2 in the liver and gonadal WAT of LLC mice **(A, B)** in the liver of fibrosarcoma (CHX)-bearing mice **(C)** and in the gonadal WAT of fibrosarcoma (MN/MCA1)-bearing mice **(D),** assessed by Western blot. Data are presented as mean ± SEM. *n* = 4 mice per group in (A), *n* = 3 in (B), *n* = 4-5 in (C) and *n* =3 in (D). Differences between groups were analyzed using an unpaired two-tailed Student’s *t* test. *p* < 0.05 was considered statistically significant.

**Suppl. Fig. 4:**
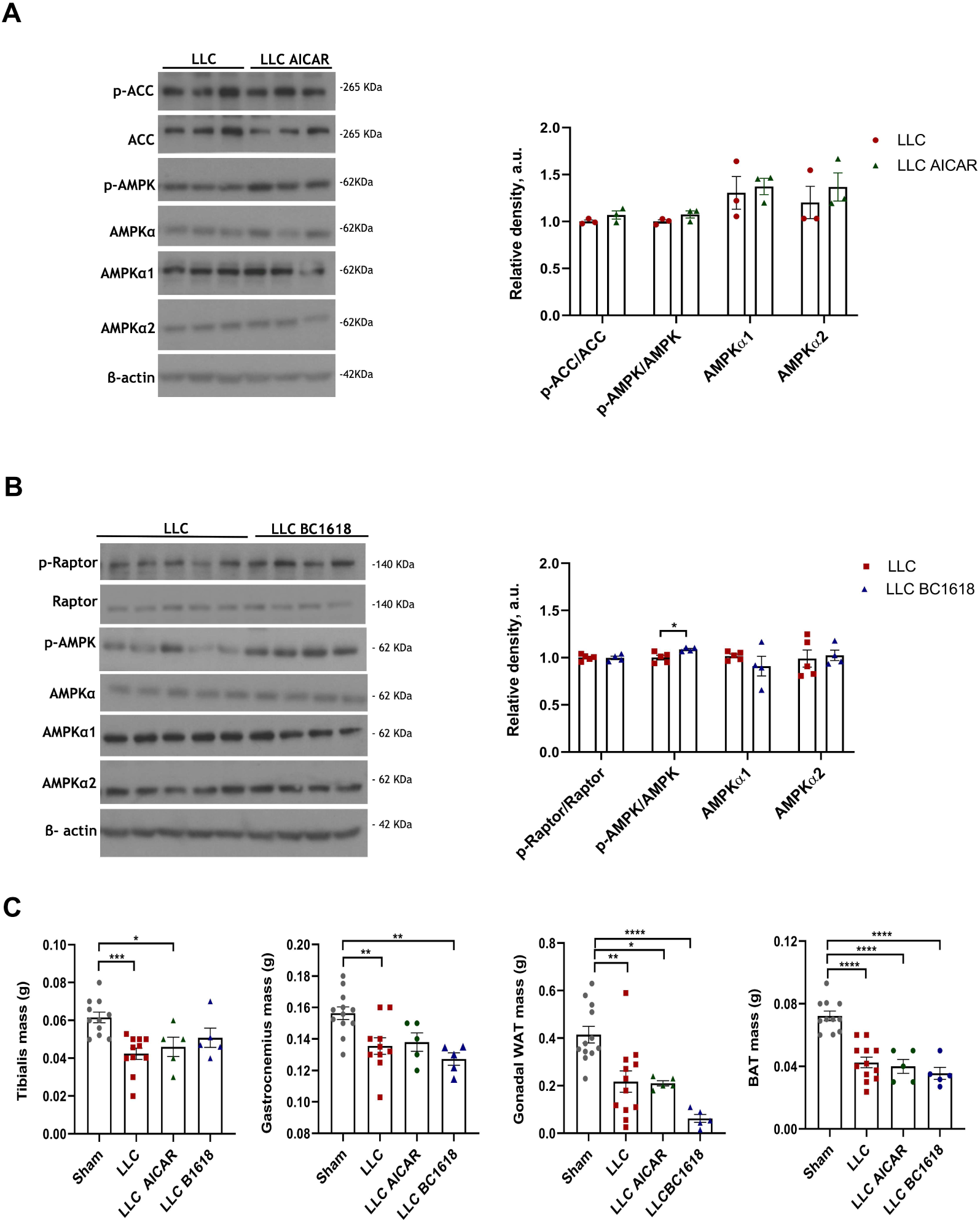
Results for Sham, LLC-bearing mice, LLC mice treated with AICAR, and LLC mice treated with BC1618 are shown. Mice were euthanized 20 days after cancer cell injection. **(A, B)** Levels of p-AMPK Thr172, total AMPKα, AMPKα1, AMPKα2, pRaptor Ser792, Raptor, ACC, pACCSer79 in the hypothalamus, determined by western blot. **(C)** Mass of tibialis muscle, gastrocnemius muscle, gonadal WAT and BAT. Data are presented as mean ± SEM. *n* = 3 mice per group in (A), *n* = 4-5 in (B) and *n* =5-12 in (C). Differences between groups were analyzed using an unpaired two-tailed Student’s *t* test (A and B) and one-way ANOVA, followed by Tukey’s test (C). *p* < 0.05 was considered statistically significant.

**Suppl. Fig. 5:**
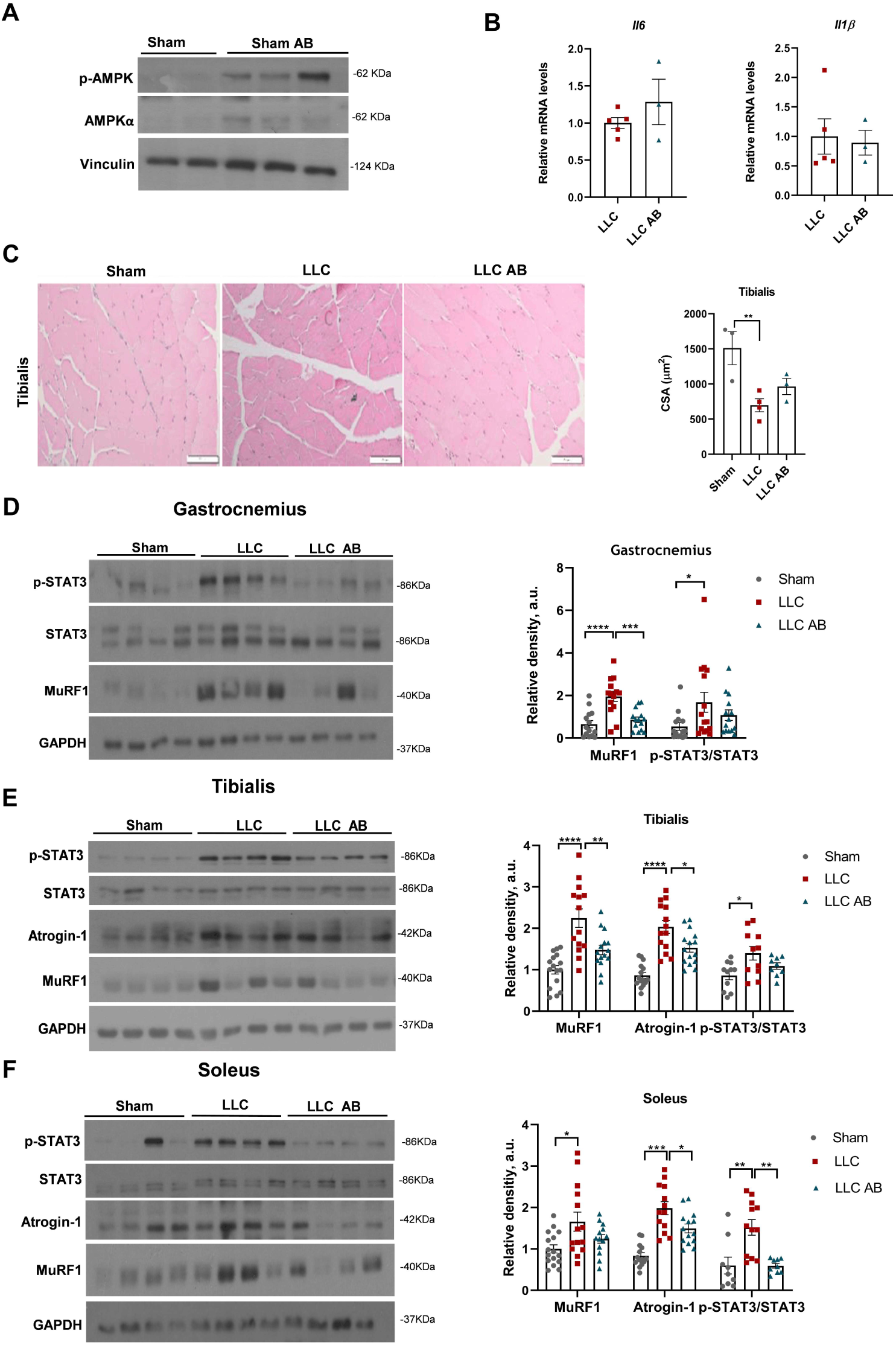
**(A)** Levels of pAMPK Thr172 and total AMPKα in Sham mice and Sham mice treated with AB, determined by Western blot. **(B)** Relative mRNA levels of *Il6* and *Il1β* in tumors from LLC-bearing mice and LLC mice treated with AB (LLC AB), determined by RT-qPCR. **(C)** Tibialis muscle fiber cross-sectional area (CSA) quantified from hematoxylin and eosin–stained histological sections. **(D-F)** Protein levels of MuRF1, atrogin1, pSTAT3 Tyr705 and STAT3 in the gastrocnemius, anterior tibialis, and soleus muscles, assessed by Western blot. Representative Western blots are shown. Data are presented as mean ± SEM. *N* = 3-5 mice per group in (B), *n* = 3 in (C) and *n* = 9-14 in (D-F). Differences between groups were analyzed using one-way ANOVA, followed by Tukey’s test. \**p* < 0.05, \*\**p* < 0.01, \*\*\**p* < 0.001, \*\*\*\**p* < 0.0001.

**Suppl. Fig. 6:**
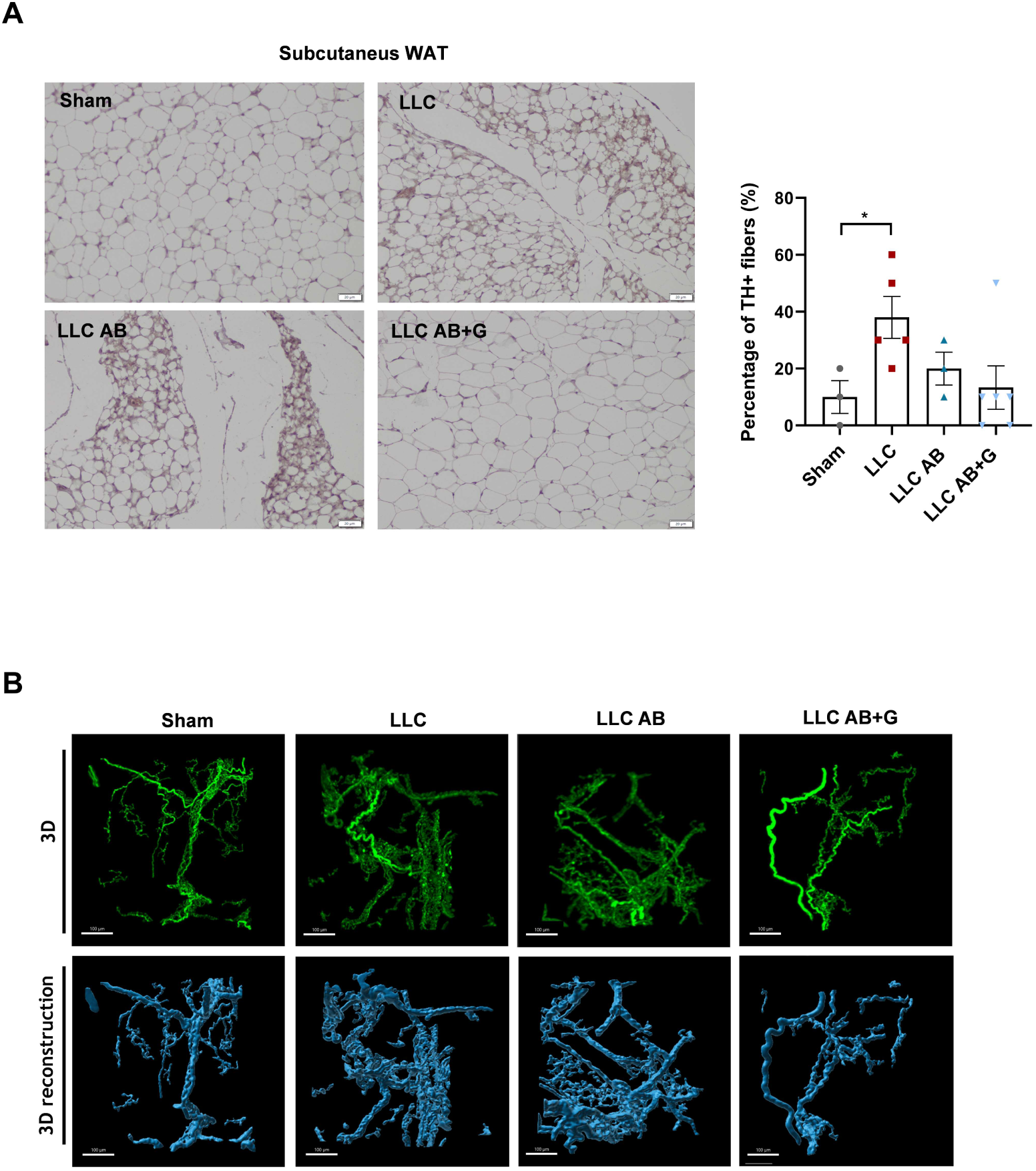
**(A)** Quantification of tyrosine hydroxylase (TH)-positive fibers (% TH) in subcutaneous adipose tissue, determined by immunohistochemical TH staining of WAT sections. *n* = 3–6 mice per group. **(B)** Representative 3D reconstruction of a selected analysis cube showing TH-positive fibers in cleared subcutaneous WAT, obtained by iDISCO and analyzed using Imaris for volumetric quantification.

