## Supplemental tables and supplemental material and methods for "Combined AMPK activation and ghrelin ameliorate cancer cachexia through complementary effects on energy homeostasis, inflammation, and wasting"

**Table S1**. List of ELISA kits used in this study

| Analyte | Kit | Reference |
| --- | --- | --- |
| **IL-6** | Mouse IL-6 Quantikine ELISA | R&D System (M6000B) |
| **IL-1β** | Mouse IL-1 beta/IL-1F2 Quantikine HS ELISA | R&D System (MHSLB00) |
| **IFN-γ** | Mouse Cytokine Milliplex | Millipore (MCYT0MAG-70K) |
| **TNF-α** | Mouse TNF Alpha | Thermo Fisher (BMS607-3) |
| **Leptin** | Mouse Leptine Quantikine ELISA | R&D System (MOB00) |
| **Corticosterone** | Corticosterone ELISA | IBL International (RE52211) |

**Table S2**. List of primers

| | Forward 5'-3' | Reverse 5'-3' |
|---|---|---|
| ***Agrp*** | TTACCCAACCTGGGCAGAAC | ATCCATTGGCTAGGTGCGAC |
| ***Npy*** | TGGACTGACCCTCGCTCTAT | GGGGCGTTTTCTGTGCTTTC |
| ***Pomc*** | GGACCTCACCACGGAGAGAGC | AGCGGAAGTGCTCCATGGAGT |
| ***Crh*** | TCTGCAGAGGCAGCAGTGCGGG | CGGATCCCCTGCTGAGCAGGGC |
| ***Murf1*** | ACGAGAAGAAGAGCGAGC | CTTGGCACTTGAGAGGAA |
| ***Atrogin1*** | ATGCACACTGGTGCAGAGAG | TGTAAGCACACAGGCAGGTC |
| ***Mhc*** | CTCTTCCCGCTTTGGTAAGTT | CAGGAGCATTTCGATTAGATCCG |
| ***Myog*** | GCAGGCTCAAGAAAGTGAATGA | TAGGCGCTCAATGTACTGGAT |
| ***Atgl*** | TCGTGTTTCAGACGGAGAGAA | CAGACATTGGCCTGGATGAG |
| ***Hsl*** | GGGTGATGAAGGACTCACCG | GATGGCAGGTGTGAACTGGA |
| ***Prdm16*** | GGAGGAGAGAGATTCCGCA | ACGTCACCGTCACTTTTGGC |
| ***Ucp1*** | CACGGGGACCTACAATGCTT | ACAGTAAATGGCAGGGGACG |
| ***Cidea*** | TGCTCTTCTGTATCGCCCAGT | GCCGTGTTAAGGAATCTGCTG |
| ***Il1β*** | GTGTCTTTCCCGTGGACCTT | AATGGGAACGTCACACACCA |
| ***Tnf-α*** | AGCCCACGTCGTAGCAAA | ATCGGCTGGCACCACTAGTTGGT |
| ***Il6*** | AGAGACTTCCATCCAGTTGCC | CCGGACTTGTGAAGTAGGGAA |
| ***Pgc1α*** | CCCTGCCATTGTTAAGACC | TGCTGCTGTTCCTGTTTTC |
| ***Hprt*** | CAGTCCCAGGGTCGTGATTA | AGCAAGTCTTTCAGTCCTGTC |
| ***Gapdh*** | CCTTGACTGTGCCGTTGAATTT | GCAAAGTGGAGATTGTTGCCAT |
| ***Slc6a4*** | TGAGCTCTTGTGTCTGTCCAT | ATAGTTTGTAATGGGCCCGGA |
| ***Tph2*** | ACCTGCGCAGTGATTTGAACA | CCCCAAGAGCTCATGCTGACA |

**Table S3**. List of antibodies

| ANTIBODIES | SOURCE | DILUTION | IDENTIFIER |
|---|---|---|---|
| **AMPKα total** | Rabbit polyclonal | 1:1000 | # 2532 Cell Signaling |
| **AMPKα1** | Rabbit polyclonal | 1:1000 | #2795 Cell Signaling |
| **AMPKα2** | Rabbit polyclonal | 1:1000 | #2757 Cell Signaling |
| **phospho-AMPKα (Thr172) (40H9)** | Rabbit polyclonal | 1:1000 | #2535 Cell Signaling |
| **Acetyl CoA Carboxylase** | Rabbit polyclonal | 1:1000 | #3662 Cell Signaling |
| **Phosho-Acetyl-CoA Carboxylase (Ser79)** | Rabbit polyclonal | 1:1000 | #3661 Cell Signaling |
| **STAT3** | Mouse monoclonal | 1:250 | ab50761 abcam |
| **phospho-STAT3(Tyr705) (D3A7)** | Rabbit monoclonal | 1:1000 | #9145 Cell Signaling |
| **HSL** | Rabbit polyclonal | 1:1000 | #4107 Cell Signaling |
| **Vinculina** | Mouse monoclonal | 1:2000 | V5405 Sigma Aldrich |
| **Phospho-HSL (Ser660)** | Rabbit polyclonal | 1:1000 | #45804 Cell Signaling |
| **ATGL** | Rabbit polyclonal | 1:1000 | #2138 Cell Signaling |
| **UCP1** | Rabbit polyclonal | 1:1000 | ab10983 abcam |
| **Raptor(24C12)** | Rabbit monoclonal | 1:1000 | #2280 Cell Signaling |
| **p-Raptor (S792) (E4v6c)** | Rabbit | 1:1000 | 89146 Cell Signaling |
| **MuRF1 (C-11)** | Mouse monoclonal | 1:500 | sc-398608 Santa Cruz |
| **MAFbx (F-9)** | Mouse monoclonal | 1:500 | sc-166806 Santa Cruz |
| **TH** | Mouse monoclonal | 1:2000 | AB1542 Millipore |
| **β-actina** | Mouse monoclonal | 1:1000 | sc-47778 Santa Cruz |
| **GAPDH (0411)** | Mouse monoclonal | 1:10000 | sc-47724 Santa Cruz |

**Table S4**. Serum levels of cytokines and hormones in sham and LLC mice

| Serum levels | Sham | LLC |
|---|---|---|
| **IL-6 (pg/ml) (*n*= 23 / 26)** | **11.41 ± 1.81** | **31.16 ± 2.89*****|
| **TNF-α (pg/ml) (*n*= 4 / 5)** | **0.24 ± 0.01** | **0.23 ± 0.015** |
| **IL-1β (pg/ml) (*n*= 6 / 6)** | **2.3 ± 0.73** | **5.85 ± 2.52** |
| **INF-γ (pg/ml) (*n*= 3 / 3)** | **2.07 ± 0.83** | **3.26 ± 1.05** |
| **Leptin (pg/ml) (*n*= 20 / 21)** | **3958 ± 620** | **806 ± 76.53*****|
| **Corticosterone (nmol/l) (*n*=23 / 23)** | **133.9 ± 8.62** | **257.8 ± 30.59 *****|

Parameters were assessed at the study endpoint. Data are presented as mean ± SEM. The number of animals per group (*n*) is shown in parentheses for Sham and LLC mice. Statistical comparisons were performed using Student's t-test. ****p*<0.001

**Table S5**. Relative mRNA expression (2^-ΔΔCt) expressed as fold change relative to Sham

| Gene | Sham | LLC | *p*-value |
|---|---|---|---|
| ***Murf1*** **(Gastrocnemius) (*n*= 6 / 5)** | **1.000 ± 0.09** | **2.493 ± 0.69** | **0.042 *** |
| ***Atrogin-1*** **(Gastrocnemius) (*n*= 6 / 5)** | **1.000 ± 0.17** | **11.50 ± 3.53** | **0.0095 **** |
| ***Mhc*** **(Gastrocnemius) (*n*= 9 / 8)** | **1.098 ± 0.15** | **0.554 ± 0.08** | **0.008**** |
| ***Myog*** **(Gastrocnemius) (*n*= 11 / 7)** | **1.000 ± 0.11** | **0.805 ± 0.06** | **0.23** |
| ***Pgc1α*** **(Gastrocnemius) (*n*= 9 / 9)** | **1.000 ± 0.21** | **0.633 ± 0.08** | **0.13** |
| ***Il6*** **(Gastrocnemius) (*n*= 4 / 4)** | **1.000 ± 0.10** | **3.278 ± 0.64** | **0.0006***** |
| ***Il1β*** **(Gastrocnemius) (*n*= 3 / 4)** | **1.000 ± 0.33** | **14.81 ± 3.43** | **0.015*** |
| ***Atgl*** **(gonadal WAT) (*n*= 9 / 8)** | **1.006 ± 0.17** | **2.353 ± 0.48** | **0.014 *** |
| ***Hsl*** **(gonadal WAT) (*n*= 9 / 8)** | **0.987 ± 0.18** | **1.419 ± 0.27** | **0.13** |
| ***Atgl*** **(subc.WAT) (*n*= 17 / 17)** | **1.000 ± 0.13** | **1.618 ± 0.36** | **0.026 *** |
| ***Hsl*** **(subc.WAT) (*n*= 17 / 17)** | **1.064 ± 0.14** | **1.360 ± 0.25** | **0.30** |
| ***Il6*** **(subc.WAT) (*n*= 5 / 5)** | **1.013 ± 0.07** | **1.746 ± 0.22** | **0.014*** |
| ***Il1β*** **(subc.WAT) (*n*= 4 / 4)** | **1.000 ± 0.108** | **24.46 ± 10.93** | **0.0472*** |
| ***Ucp1*** **(subc.WAT) (*n*= 15 / 17)** | **1.000 ± 0.30** | **2.398 ± 1.15** | **0.27** |
| ***Cidea*** **(subc.WAT) (*n*= 14 / 17)** | **0.980 ± 0.15** | **1.750 ± 0.56** | **0.14** |
| ***Prdm16*** **(subc.WAT) (*n*= 4 / 5)** | **1.000 ± 0.08** | **0.787 ± 0.20** | **0.39** |
| ***Ppargc1α*** **(subc.WAT) (*n*= 9 / 7)** | **1.000 ± 0.18** | **1.003 ± 0.20** | **0.99** |

Parameters were assessed at the study endpoint. Data are expressed as mean ± SEM. The number of animals per group (*n*) is indicated in parentheses for Sham and LLC mice. Statistical comparisons were performed using Student's t-test. *$p<0.05$, **$p<0.01$, ***$p<0.001$, ****$p<0.0001$

### Supplemental methods

#### Additional cachexia mouse models

To validate selected findings, tissue samples obtained from additional cachexia mouse models were analysed. The mice fibrosarcoma CHX model was generated in the laboratory of Prof. M. Schweiger and Prof. R. Zechner (Institute of Molecular Biosciences. University of Graz. Austria) (Ref. S1). Male C57BL/6 mice were injected with 1 × $10^6$ CHX 207 fibrosarcoma cells into the gastrocnemius muscle of the right hind limb. Mice were euthanized 16 days after cancer cell implantation. Hypothalamus, liver, and white adipose tissue samples from these mice were examined.

In addition, white adipose tissue (WAT) from female mice bearing MN/MCA1 fibrosarcoma tumors were evaluated. In this model, 1 × $10^5$ MN/MCA1 cells were injected into the caudal thigh muscle, and animals were sacrificed 25 days after tumor cell inoculation.

Furthermore, hypothalamus from chronic Lymphocytic Choriomeningitis virus (LCMV)-infected C57BL6/J mice were studied. These tissues were kindly provided by Prof. R. Zechner and A. Bergthaler from Institute of Molecular Biosciences. University of Graz and CeMM Research Center for Molecular Medicine of the Austrian Academy of Sciences, Vienna, Austria (Ref. S2). In this model, wild-type C57BL/6J mice were infected with 2 × $10^6$ focus-forming units (FFU) of the chronic Clone 13 strain of LCMV and sacrificed 12 days post-infection.

#### cDNA synthesis and qPCR

Total RNA was extracted from frozen tissue using TRIzol reagent (Invitrogen, Carlsbad, CA) according to the manufacturer's instructions. cDNA was synthesized using a mix of MgCl2 (2.5 mM), dNTPs (0.5 mM), random primers (17 ng/µL), RNase OUT ribonuclease inhibitor (33.3 U/mL), and M-MLV reverse transcriptase (13.3 U/mL) in 5× First Strand Buffer. Real-time PCR was performed using the Luminaris Color HiGreen qPCR Master Mix Kit (ThermoFisher Scientific, USA) and QuantStudio V.7 Flex Real Time PCR Systems (Applied Biosystems) in a total reaction volume of 10 µL. Cycling conditions included an initial denaturation at 95ºC for 10 minutes followed by 40 cycles at 95ºC for 15 seconds and 60ºC for 1 minute, with a holding stage of 95ºC for 15 seconds, 60ºC for 1 minute and 95 ºC for 15 seconds. All reactions were performed in triplicate. Gene expression was normalized relative to the housekeeping gene GAPDH (for muscle and fat) or HPRT (for hypothalamus) and results were expressed as fold change to the control using the 2−ΔΔCt. All primer sequences are shown in **Table S2**.

#### Western Blot Analysis

Proteins were extracted from frozen tissues and were lysed with lysis buffer for hypothalamic samples and with RIPA for muscle and adipose tissue**.** Samples were homogenized in the Tissue Lyser II (Qiagen) at 30 Hz for 3 minutes. Lysates were centrifuged at 13,000 rpm for 15 minutes at 4 °C, and protein concentration was measured using the Bradford microassay (Bio-Rad Laboratories, CA, USA) when lysis buffer was used or the BCA assay for RIPA samples. In both cases, the manufacturer's instructions were followed. Protein samples were separated by SDS-PAGE in 8%-12% acrylamide gels using a Mini-PROTEAN Tetra Cell electrophoresis system (Bio-Rad Laboratories, CA, USA) and blotted onto polyvinylidene difluoride (PVDF) membranes (BioRad) using a Bio-Rad Trans-blot SD Semi-dry Transfer Cell (Bio-Rad Laboratories, CA, USA). Membranes were blocked with 5% BSA in TBS-Tween 0,2% for 2 h at room temperature and incubated with the primary antibodies listed **Table S3** overnight at 4 ºC. Membranes were washed with TBS-Tween 0,2% for 5 min 4 times and incubated with the appropriate horseradish peroxidase-conjugated secondary antibodies (1:5000) for 1h at room temperature. After washing the membranes 4 times to remove excess secondary antibody, the Pierce ECL Western chemiluminescent substrate (Thermo Fisher Scientific, MA, USA) was applied. The chemiluminescent signals were detected by exposing the membranes to photosensitive films (FUJI Medical X-Ray Film), which were developed in a Curix 60 processor (AGFA Healthcare). The films were digitized, and band intensities were quantified by densitometric analysis with ImageJ (http://rsbweb.nih.gov/ij). Protein levels were expressed in relation to β-actin (for hypothalamus), GAPDH (for muscle) and vinculin (for WAT) protein levels.

**Skeletal muscle and adipose tissue histology.**

For histopathological analysis, skeletal muscle and adipose tissue samples were fixed for 24 hours in 10% formalin buffer and then were dehydrated and embedded in paraffin following standard procedures. Sections of 4 µm were cut using a microtome and stained with an alcoholic Hematoxilin/Eosin (H&E) protocol (BioOptica) following the manufacturer's instructions. Histological sections were examined using an Olympus IX73 microscope (20x objective) equipped with an Olympus DP74 camera. To quantify the cross-sectional area (CSA) of tibialis and gastrocnemius muscle or the sizes of adipocytes in subcutaneous fat, three to four fields per sample were randomly chosen and analyzed using ImageJ (https://imagej.nih.gov/ij/). Values were expressed as the mean for each experimental group.

**Immunohistochemistry**

For immunohistochemistry, paraffin sections were deparaffinized with xylene and rehydrated in a graded ethanol series. After antigen retrieval, sections were incubated overnight at 4ºC with TH primary antibody (1:1000, Millipore, AB1542, Temecula, CA, USA). After several washes, the sections were incubated with anti-goat biotinylated secondary antibody (EnVision System HRP, Agilent DAKO, Glostrup, Denmark) for 1 h at 37 ºC. To visualize peroxidase activity, 3,3'-diaminobenzidine (DAB; OriGene, Beijing, China) was used as a chromogen. Images were observed using a light microscope (Olympus IX73) and quantified using ImageJ (https://imagej.nih.gov/ij/).

**iDISCO** (Figure SM1)

*Perfusion and tissue processing*

Mice were anesthetized with a mixture of ketamine and xylazine injected intraperitoneally. Tissues were fixed by intracardiac perfusion with 4% paraformaldehyde in phosphate-buffered saline (PBS). Subcutaneous white adipose tissue samples were carefully dissected and post-fixed in 4% paraformaldehyde at 4°C overnight. The samples were stored in PBS at 4°C until further processing.

*Dehydration and Delipidation*

The samples were dehydrated in an increasing graded methanol series (20%, 40%, 60%, 80%, and 100% in water) for 1 hour per wash. To remove residual water, an additional 2-hour wash was performed in 100% methanol with agitation at room temperature. The samples were then delipidated overnight with agitation at room temperature in a solution of 66% dichloromethane (Sigma-Aldrich, St. Louis, MO, USA) in methanol. Following delipidation, the samples were washed twice in 100% methanol (4 hours per wash) to remove remaining dichloromethane.

*Tissue bleaching*

Samples were bleached with a solution of methanol containing a 5% of hydrogen peroxide (Sigma-Aldrich, St. Louis, MO, USA) overnight at 4ºC in the dark without shaking.

*Rehydration and permeabilization*

The samples were rehydrated in a graded methanol series (60%, 40%, and 20%) for 1 hour per wash at room temperature with agitation. They were then washed twice in PBS for 15 minutes each, followed by a third 15-minute wash in PBS containing 0.2% Triton X-100 (Sigma-Aldrich, St. Louis, MO, USA). Finally, the samples were permeabilized via a 24-hour incubation at 37 °C in a solution of 20% dimethyl sulfoxide (Sigma-Aldrich, USA) and 2.3% glycine (Sigma-Aldrich, St. Louis, MO, USA) in PBS-T (PBS with 0.2% Triton X-100).

*Immunolabeling*

Prior to immunostaining, samples were blocked in PBS-T containing 0.2% gelatin (Sigma-Aldrich, St. Louis, MO, USA) for 24 hours at 37 °C; this same blocking buffer was used for all subsequent antibody dilutions. To label UCP1 and the sympathetic innervation of the adipose tissue, samples were incubated with a combination of primary antibodies against UCP1 (1:1000, Abcam, ab10983) and tyrosine hydroxylase (TH; 1:2000, Millipore, AB1542, Temecula, CA, USA) for 10 days at 37 °C with shaking. Post-incubation, the samples were washed four times over two days in PBS containing 0.2% Tween 20 and 1 U/mL heparin (PBS-TwH). The samples were then incubated for 10 days in secondary antibodies diluted in blocking buffer: Alexa Fluor 568-conjugated secondary antibody (1:1000, Invitrogen) to detect UCP1, and Alexa Fluor 647-conjugated secondary antibody (1:1000, Invitrogen, Carlsbad, CA, USA) to visualize sympathetic innervation. Finally, the samples were washed four times over two days in PBS-TwH at room temperature with shaking in the dark.

*Tissue Clearing*

For ease of handling under the microscope, the samples were embedded in 1% agarose in water. They were then dehydrated in an increasing graded methanol series (20%, 40%, 60%, and 80% for 1 hour each, followed by 100% methanol overnight). Each step was carried out at room temperature with agitation and in the dark. Following dehydration, samples were incubated in a solution of 66% dichloromethane and 33% methanol for 3 hours. Residual methanol was removed with two final 15-minute washes in 100% dichloromethane. Finally, the samples were cleared in dibenzyl ether (DBE; Sigma-Aldrich, St. Louis, MO,

USA) overnight at room temperature in the dark without shaking. Samples were stored under these conditions until light-sheet imaging.

*3D Imaging*

All whole-tissue samples were imaged on a light-sheet microscope (Blaze, Miltenyi BioTec) equipped with an sCMOS camera and 1.3× and 4× objective lenses, used for low- and high-magnification whole-tissue views, respectively. Images were acquired using ImspectorPro software (Miltenyi BioTec). Samples were illuminated bi-directionally and scanned using laser lines at 488, 561, and 640 nm. Optical sectioning spacing was set to 6 µm for the 1.3× objective and 3 µm for the 4× objective

*Image Processing*

All whole-tissue images were analyzed and visualized using Imaris x64 software (version 10.2, Bitplane). To generate high-resolution tile scans, multiple images acquired at 4× magnification were stitched using the Imaris Stitcher module (version 10.2, Bitplane). Three-dimensional (3D) reconstructions were then generated using the 'volume rendering' function.

*Quantification of Sympathetic Innervation Volume*

Within each subcutaneous WAT sample, five small cubic regions located within UCP1-positive areas were randomly selected **(Figure** SM2). These cubes were generated using the Imaris Surface tool. The image was segmented to create a 3D surface corresponding to the signal of interest (TH+ fibers). A mask was then applied to remove the background signal. The Volume function within the Surface tool was used to reconstruct and quantify the volume of TH-positive fibers within each cube (pixels$^{3)}$ **(Figure** SM2**).** Finally, the mean of TH-positive fiber volume across the five cubes was calculated to obtain a single value for each animal.

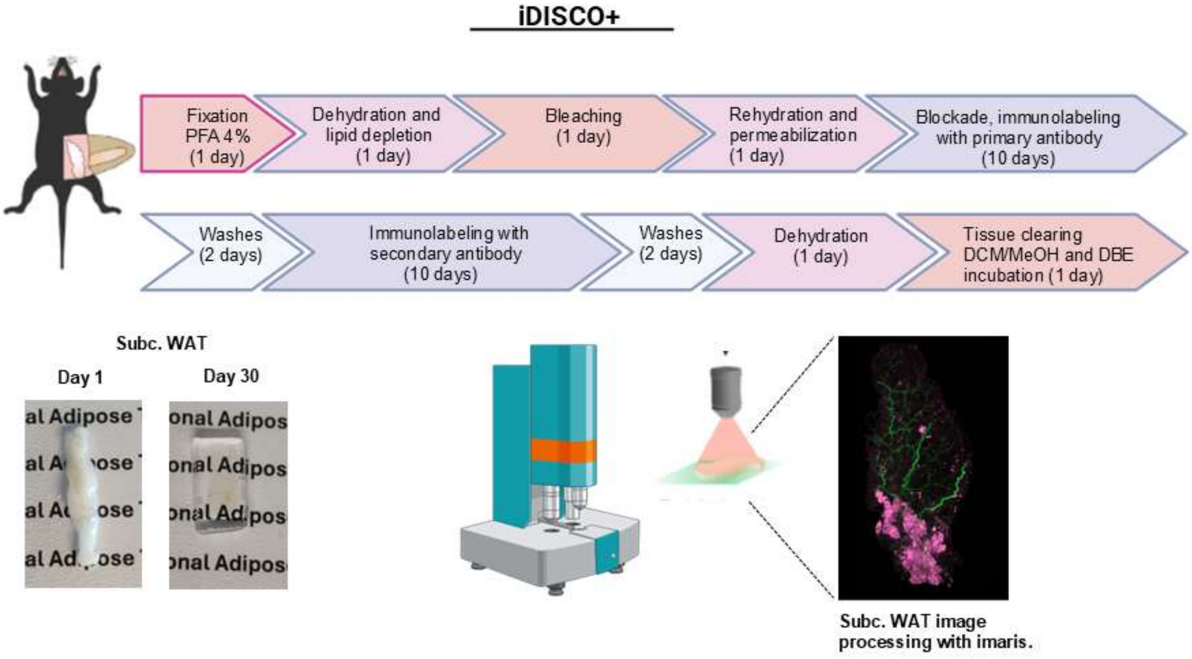


Figure Suppl. Methods 1

Figure Suppl. Methods 2

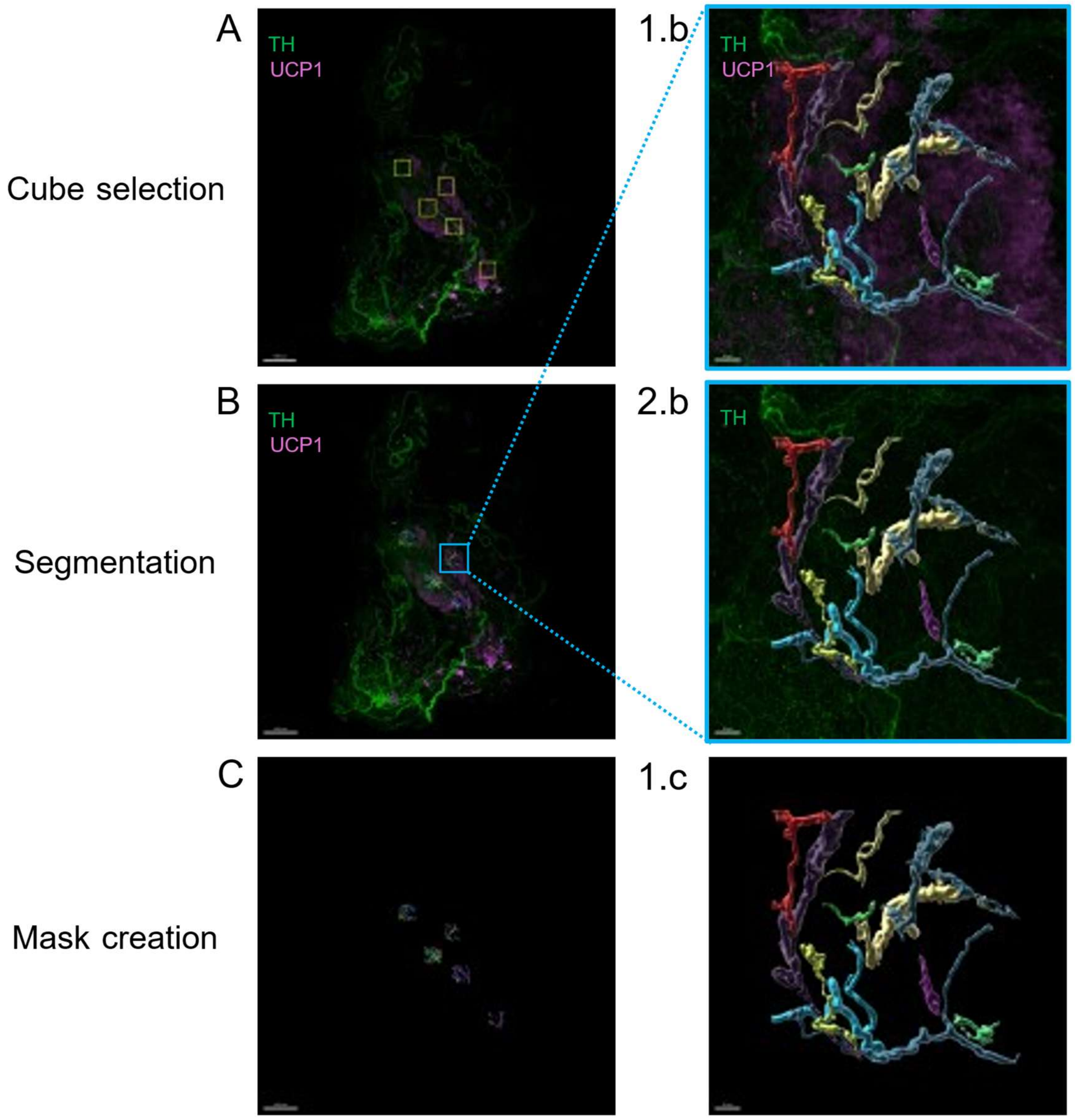
